# Sound-Evoked Plasticity Differentiates Tinnitus from Non-Tinnitus Mice

**DOI:** 10.1101/2024.12.20.629689

**Authors:** Emily M. Fabrizio-Stover, Christopher M. Lee, Douglas L. Oliver, Alice L. Burghard

**Affiliations:** Department of Otolaryngology, Medical University of South Carolina, Charleston, SC, 29414, United States of America; Department of Neuroscience, University of Connecticut School of Medicine, Farmington, CT, 06030 United States of America

**Keywords:** tinnitus, inferior colliculus, auditory brainstem response, sound-evoked plasticity, noise exposure

## Abstract

Tinnitus is the perception of non-meaningful sound in the absence of external stimuli. Although tinnitus behavior in animal models is associated with altered central nervous system activity, it is not currently possible to identify tinnitus using neuronal activity alone. In the mouse inferior colliculus (IC), a subpopulation of neurons demonstrates a sustained increase in spontaneous activity after a long-duration sound (LDS). Here, we use the “*LDS test*” to reveal tinnitus-specific differences in sound-evoked plasticity through IC extracellular recordings and the auditory brainstem response (ABR_LDS_) in CBA/CaJ mice after sound exposure and behavioral tinnitus assessment. Sound-exposed mice showed stronger and shorter tone-evoked responses in the IC compared to unexposed controls, but these differences were not strong predictors of tinnitus. In contrast, in the LDS test, non-tinnitus mice had a significantly stronger suppression in tone-evoked spike rate compared to tinnitus and unexposed control mice. ABR peak amplitudes also revealed robust differences between tinnitus and non-tinnitus mice, with ABR peaks from non-tinnitus mice exhibiting significantly stronger suppression in the LDS test compared to tinnitus and control mice. No significant differences were seen between cohorts in ABR amplitude, latency, wave V:I ratio, wave V:III ratio, I-V intra-peak latency, and I-VI intra-peak latency. We found high-frequency tone stimuli better suited to reveal tinnitus-specific differences for both extracellular IC and ABR recordings. We successfully used the LDS test to demonstrate that tinnitus-specific differences in sound-evoked plasticity can be shown using both multi-unit near-field recordings in the IC and non-invasive far-field recordings, providing a foundation for future electrophysiological research into the causes and treatment of tinnitus.

## 1 Introduction

Evidence of tinnitus behavior in animal models of tinnitus is associated with increased spontaneous firing rate (SFR) at many levels of the auditory nervous system (Kalappa et al. 2014, Brozoski et al. 2002, Kaltenbach et al. 2005, Sametsky et al. 2015), including the inferior colliculus (IC) (Bauer et al. 2008, Longenecker and Galazyuk 2011, Holt et al. 2010). However, there is little information about sound-driven responses in the IC or related sound-evoked plasticity in animals with behavioral evidence of noise-induced tinnitus.

A *long-duration sound test* (LDS test) developed in the Oliver Laboratory, consisting of recordings before and after a *long-duration sound* (LDS), has been shown to detect sound-induced plasticity in the central nucleus of the IC (ICC) of wild-type mice (Burghard 2022). In extracellular multi-channel electrode recordings from the ICC, roughly 16% of channels had a sustained increase in spontaneous activity, referred to as a *long-duration sound-induced afterdischarge* in the silent period after the LDS. Furthermore, approximately 16% of the total responsive channels had facilitated responses to sound after an LDS. Thus, the LDS altered both the spontaneous activity and sound-driven response in the ICC.

If neurons that generate an afterdischarge in the ICC become chronically hyperactive in noise-induced tinnitus, that hyperactivity may generate an identifiable electrophysiological signal indicating tinnitus. The auditory brainstem response (ABR) is a recording method representing the synchronized neural activity along multiple points in the auditory brainstem, including the IC (Melcher and Kiang 1996). ABRs have been unsuccessful in distinguishing between tinnitus and non-tinnitus patients and animals with and without evidence of tinnitus behavior (Domarecka et al. 2020, Milloy et al. 2017, Jacxsens et al. 2022). However, if the LDS is shown to generate tinnitus-specific changes in sound-driven activity in the ICC, and because the IC contributes to the ABR (Land et al. 2016), then differences in sound-driven activity between tinnitus and non-tinnitus subjects could be seen in the ABR.

Here, the LDS test was used to investigate changes in sound-evoked plasticity in mice with and without evidence of tinnitus behavior. Two types of electrophysiological recordings were used to examine how the LDS test affects the central auditory system. We recorded multi-unit near-field activity in the ICC using multi-channel electrodes and far-field activity with ABR_LDS_ recordings. All recordings were compared in mice with behavioral evidence of tinnitus after sound exposure, mice without behavioral evidence of tinnitus after sound exposure, and in control, unexposed mice. Tinnitus status was determined using active avoidance behavior. Whenever possible, the same mice were used for both ABR_LDS_ and IC recordings, thus allowing a comparison of the LDS-induced changes in both recording types for mice with tinnitus, no tinnitus, or no sound exposure history. We found that with both methods, mice with sound exposure but no behavioral sign of tinnitus showed a reduced response to sound after the presentation of an LDS. In contrast, animals with behavioral signs of tinnitus responded similar to the control (not sound-exposed) mice.

## 2 Materials and methods

### Animals

This animal study was approved by local Institutional Animal Care and Use Committee (IACUC). Experiments were performed on CBA/CaJ mice (Jackson Laboratories; Strain #000654, RRID: IMSR_JAX:000654, Bar Habor, ME, USA) of both sexes. A total of 68 CBA/CaJ mice were used in this study. All mice were purchased at the age of 4-8 weeks and housed five in a cage employing a 12-hour light/dark cycle with continuous access to food and water. Additional nesting materials were added as enrichment. All experiments were performed in accordance with institutional guidelines and the NIH Guide for the Care and Use of Laboratory Animals and were approved by the Animal Care and Use Committee at the University of Connecticut Health Center.

All animals underwent a hearing assessment, and the experimental animals underwent a behavioral training phase. Control animals had no behavioral training. After successfully learning the behavioral paradigm, experimental animals were exposed to loud sound, and hearing was re-assessed 2-4 weeks later. Eight weeks after the sound exposure, the experimental animals were re-tested in the behavioral paradigm. Following this, the experimental and control animals underwent an ABR recording before and after an LDS (ABR_LDS_) presentation. Later, they underwent surgery, and IC multi-unit recordings were performed in response to the LDS test stimulus. All electrophysiological recordings were done under general anesthesia.

### Anesthesia

For hearing assessments before and after sound exposure as well as the ABR_LDS_ recordings, animals were anesthetized using a mixture of ketamine and xylazine in saline (ketamine 100 mg/kg, xylazine 14.3 mg/kg) administered *i.m.* Anesthesia was maintained by alternating injections of 50 mg/kg ketamine and a mix of 50 mg/kg ketamine and 7.1 mg/kg xylazine. A maximum volume of 0.1ml per 10 grams of mouse body weight was used for both solutions. Injections were performed in the hindlimb of the mouse or *i.p*. For the IC recordings, induction was the same, but anesthesia maintenance was done via isoflurane 0.5-2 % in 100 % oxygen; otherwise, anesthesia monitoring was the same for all procedures. After induction, animals were placed in a gas anesthesia head holder (David Kopf Instruments, Tujunga, CA, USA), which provided 100 % oxygen (0.5 L flow rate). Anesthesia depth was checked every ∼30 min via toe pinch reflex, and heart rate, breath rate, and blood oxygenation were constantly monitored via pulse oximetry (Mouse Ox, Starr Life Science Corp, Oakmont, PA, USA). Body temperature was maintained at 36-37° C by placing the animal on a heating pad coupled with a rectal thermometer.

To briefly anesthetize the animals and introduce the earplug into the pinna for unilateral sound exposure, they were placed in an induction chamber and exposed to 4 % isoflurane in 100 % oxygen at a flow rate of 2 L/min until they lost consciousness.

### Surgery

The surgery necessary for the IC recordings was like the one described in detail in Burghard et al. (2022). In short, the skull over the IC (bilaterally in the current study) was removed using a dental drill. Following bone removal, a hole was drilled in the skull over the left parietal lobe to place a screw that served as a reference electrode. After the screw was anchored in place, the dura mater was removed to expose the ICs. Ice-cold saline was used to keep the brain surface moist and to stop any potential bleeding.

### Hearing assessment (before and after sound exposure)

Using click-evoked ABR and amplitude modulation following response (AMFR) measurements, the hearing status of all mice (except control mice) was assessed before any further testing or behavioral training. Click-evoked ABRs were used to identify absolute hearing thresholds and AMFRs were used to determine frequency-specific thresholds. Those recordings were performed on anesthetized animals. The methods for collecting click-evoked ABR and AMFR were published previously (Burghard et al. 2019). Briefly, needle electrodes (Genuine Grass Reusable Subdermal Needle Electrodes, Natus, Middletown, WI, USA) were inserted under the skin, behind each ear, and at the vertex of the head. In the current study, foam earplugs (CVS Health Foam Earplugs Advanced Protection, 30-decibel reduction rating, CVS Health Corporation, Woonsocket, RI, USA) were used to help isolate responses from individual ears.

All recordings were performed in a sound-attenuated chamber (IAC, Bronx, NY, USA). Sounds were presented via a calibrated free-field speaker (Revelator R2904/7000-05 Tweeter, ScanSpeak, Videbæk, Denmark). An RZ6 Acoustic Processor generated acoustic stimuli (Tucker Davis Technologies, TDT, Alachua, FL, USA), and responses were digitized at a sampling rate of 25 kHz using a TDT RA4L1 head stage. BioSig software (TDT) was used to evoke and analyze ABRs. Click stimuli were presented at a presentation rate of 21 Hz and in ascending 5 dB steps. Hearing thresholds were determined by the midpoint between the stimulus intensity of the first detectable ABR waveform and the last stimulus intensity without a detectable waveform.

The AMFR procedure was performed as described in Burghard et al. (2019). Custom code (©Gongqiang Yu, UConn Health) in MATLAB (MathWorks, Natick, MA, USA) and TDT RPvdsEX generated the acoustic stimuli and collected and analyzed the AMFR. The AMFR was evoked with continuous, amplitude-modulated, 1/3 octave bandpass-filtered noises centered at 8, 11, 16, 22, or 32 kHz. A modulation frequency of 42.9 Hz was chosen to focus on auditory brainstem generators of the AMFR response (Kuwada et al., 2002). Stimuli were presented in descending 10 dB steps. The starting dB SPL of the AMFR was ∼30 dB above the previously determined click ABR threshold. The AMFR threshold was identified as the midpoint between the lowest dB SPL level with a response and the highest dB SPL level without a response.

### Sound Exposure

The procedure to expose mice to loud, damaging acoustic stimuli has been described previously (Fabrizio-Stover et al. 2022), and the term “sound-exposed” (SE) is used here to refer to these mice regardless of whether they developed tinnitus. A foam earplug was inserted into the right ear canal and held in place with Liquid Bandage (CVS Health Corporation) to help preserve normal hearing in that ear, with the left ear fully exposed to the sound. After inserting the earplug, the mouse was allowed to recover from the brief anesthesia for at least 20 minutes before sound exposure. Sound exposure was performed in an anechoic chamber (IAC Acoustics, Naperville, IL, 28’x19’x17’) using a pair of calibrated Eminence N151M 8Ω speakers (Eminence Speaker LLC, Eminence, KY, USA) modified with a Ferrofluid Retrofit Kit (Ferro Tec #020618-L11, Bedford, NH, USA) and mounted on Eminence H290B horns facing each other presenting uncorrelated narrowband noise. During the sound exposure, two mice were housed separately in two neighboring acoustically transparent holding cage compartments. Mice were exposed to a 2 kHz wide, 16 kHz-centered 113 dB SPL narrowband noise for one hour. Previously, we demonstrated that this sound exposure paradigm did not result in significantly different pure tone threshold shifts between tinnitus and non-tinnitus mice (Fabrizio-Stover et al. 2022). Mice were monitored continuously via a webcam during sound exposure and observed for signs of discomfort or distress. After sound exposure, the earplug was removed, and the mice were returned to their home cages.

To confirm that the earplug spared the right ear from trauma, bilateral or right ear unilateral hearing thresholds were reassessed with ABR and AMFR at 2-4 weeks after sound exposure. Unilateral hearing thresholds were collected with one ear plugged with a foam earplug, and auditory stimuli were presented in an open field. Animals with auditory thresholds higher than 65 dB SPL with binaural ABRs or ABRs collected from the unexposed ear were excluded from further behavioral testing (n=5).

### Behavioral Tinnitus Assessment

Behavioral tinnitus assessment was performed using the Active Avoidance (AA) method. It was based on changes in response to a conditioned stimulus developed by Dr. Bradford May (The Johns Hopkins University) and has been described in detail (Fabrizio-Stover et al. 2022). In short, mice were trained in a two-chamber shuttle box (PanLab, Harvard Apparatus, model LE916-918, Barcelona, Spain) housed in a sound-attenuated chamber. Tones (9-36 kHz, ¼ octave step size) were presented randomly to the mouse. The duration of the tones was fifteen seconds maximum, with five seconds of tone presentation before a shock would be administered. The mouse could avoid the shock and stop sound presentation if it moved to the other side of the chamber. Tone presentation would stop at fifteen seconds or when the mouse moved to the other side of the chamber, whichever occurred first. A TDT RP2 processor generated all sound stimuli.

Each session consisted of approximately 70 stimulus trials and lasted approximately 45 minutes, including a 5-minute habituation period at the start of the session. Animals underwent one training session each day during the light phase. Animals that performed at 75% avoidance accuracy over five consecutive days with 60-70 dB SPL stimulus presentation levels were used in this study. Eight animals were excluded due to failing to reach this threshold. Eight weeks after sound exposure, AA performance was again assessed over five sessions. To prevent the mice from learning to distinguish their internal percepts from external sound presentation, sessions were conducted once or twice a week on non-consecutive days. Shocks were only presented during 50% of the unsuccessful trials.

### Auditory brainstem response recordings using the LDS test (ABR_LDS_)

ABR_LDS_ recordings were made from 16 tinnitus mice, 7 non-tinnitus mice, and 5 control mice. The recording location, setup, anesthesia, equipment, recording set-up, and acoustic stimuli generation were the same as those used for the ABR and AMFR procedures. Recordings were isolated from each ear by using a piece of foam earplug to block the pinna of the ear that was not of interest. Control, unexposed mice always had their right pinna blocked, so responses were driven primarily by the left ear. The impedance of the ground and reference electrodes was less than 1 kΩ.

The LDS was a 1/3 octave, band-passed noise of 60 seconds duration, generated by applying a finite impulse response filter (FIR) (12 dB/octave roll-off) to Gaussian noise. Three LDS center frequencies were used: one below the sound exposure center frequency (<15 kHz, typically 8 kHz), one matching the center frequency of the sound exposure stimulus (16 kHz) or matching the tinnitus frequency as identified by AA in tinnitus mice, and one that was above the sound exposure center frequency (>17 kHz, typically 32 kHz) or a second tinnitus frequency if relevant. The center frequency of the ABR stimulus (3 ms tone pip, 1 ms cosine rise/fall time) before (PRE) and after (POST) the LDS matched the center frequency of the LDS. The tone pips were presented in 6 trains of 10 seconds at 21.1 Hz, separated by 10 seconds of silence. This presentation pattern was identical for the PRE and POST LDS stimuli. All stimuli were presented at least 30 dB SPL above the frequency threshold as determined with the AMFR or at the maximum stimulus level possible with the equipment used (90 dB SPL).

### Extracellular Recording in the ICC

Extracellular multi-unit recordings were made in the ICC of 14 sound-exposed mice with behavioral evidence of tinnitus (tinnitus), 8 without behavioral evidence of tinnitus (non-tinnitus), and 6 control, unexposed mice. Data were collected from the ICC ipsi-and contralateral to the noise-exposed ear (right ICC for control animals). The recording setup was the same set-up as the previous electrophysiological recordings, and the procedure was the same as in Burghard et al. (2022) with the additional recording from the second IC. Acoustic stimuli were generated with an RZ6 processor (TDT) at a sampling rate of 200 kHz. Parameters of the acoustic stimuli were set with custom MATLAB software and then transmitted to “Synapse” software (TDT) via the MATLAB function “SynapseLive” (TDT). Broadband noise bursts (3-50 kHz, 85- or 90-dB SPL, 100 ms duration, 2 Hz presentation rate) were played during electrode placement and the presence of sustained sound-driven responses confirmed the location of the electrodes in ICC. Responses were collected with custom 32-channel, 2-shank linear silicon probes (length: 3 mm, 16 channels/shank, Neuronexus, Ann Arbor, MI, USA). The impedance of the electrode sites ranged from 0.22 to 1.68 MΩ. The two shanks were spaced 400 µm apart, and the electrode sites were placed 100 µm apart. The probe was inserted with a manipulator (Scientifica, Uckfield, UK) at an angle of 10 degrees pitched caudal from the vertical. The average channel depth was 1.88 mm (±0.16 mm STD). Electrode signals were digitized at 25 kHz with a TDT PZ5 amplifier and delivered to a TDT RZ5 processor. The signals were filtered with a 30 Hz hi-pass and spikes were detected by thresholding the voltage signal. Thresholds on each channel were manually adjusted and were typically ∼5x the signal standard deviation. Frequency response areas (FRA) were obtained by presenting a random sequence of pure tones (200 ms duration, 4-64 kHz, 0-90 dB SPL, 10 dB, and 0.25 octave). Each tone/sound level combination was presented five times.

For each ICC, up to six different recordings were performed. Spontaneous activity recordings were collected before and after an LDS with three different LDS center frequencies to determine the percentage of channels with a long-duration sound induced afterdischarge. Then, for the LDS test, recordings collected sound-driven activity before and after an LDS with the same three LDS center frequencies. The LDS stimulus was the same as used in the ABR_LDS_. Three LDS center frequencies were used for each mouse. In non-tinnitus mice, these frequencies were 8 kHz, 16-21 kHz, and 32 kHz. In tinnitus mice, these frequencies were 8 kHz, the tinnitus frequency indicated in AA (usually between 16-21 kHz), and either 32 kHz or the second tinnitus frequency indicated in AA. In each IC, spontaneous and sound-driven activity would be collected before and after the LDS. Spontaneous and sound-driven responses would be collected for each pre-determined center frequency. For sound-driven responses, the frequency of the tones would be matched to the LDS. Therefore, six recordings (3 different frequencies X 2 conditions) would be collected from each IC. The LDS center frequency was presented 30 dB SPL above the lowest pure tone threshold across all channels responding to that respective frequency as determined in the FRA (minimum absolute LDS level: 60 dB SPL, maximum 90 dB SP (the maximum intensity possible with the system).

Spontaneous activity was studied by measuring activity during a 60 second silent period before the LDS and during a 240-second silent period after the LDS. Sound-driven activity in the IC was studied by using trains of tone-pips presented in 6 trains of 21.1 Hz, separated by 10 seconds of silence, with 6 trains before the LDS and 6 trains after the LDS. Extracellular recordings were collected from the ICC contralateral to the sound-exposed ear first, then from the ipsilateral ICC to the sound-exposed ear. The procedure was the same for both sides. New pure tone thresholds for the ipsilateral side were based on a second FRA recorded from this IC and used to determine stimulus level. The same LDS center and tone pip frequencies were used for recording from both ICCs. In control mice, recordings were only done from the right ICC.

### Analysis

#### Behavioral Analysis

A correct AA trial was classified as relocation of the mouse after tone onset but before shock onset (within the first 5 seconds of tone onset). Performance in AA was recorded as correct avoidance or no avoidance if the shock was not avoided. The percentage of correct avoidance responses from sound-exposed animals was averaged across five days. Mice with tinnitus are hypothesized to exhibit reduced avoidance behavior when the presented stimulus is similar or overlapping with the tinnitus. As described previously, the mean percent correct avoidance score was calculated across all tested frequencies after sound exposure (Fabrizio-Stover et al. 2022). To determine tinnitus, the frequency with the lowest average percent correct avoidance was found and compared to the average percent correct avoidance using a one-sample, one-tailed t-test. If the tested frequency with the lowest correct avoidance was significantly lower than the average correct avoidance (p<0.05), then the animal was identified as a tinnitus mouse.

#### Analysis of ABR_LDS_

The ABR_LDS_ data were analyzed with custom MATLAB (MathWorks) software (© Christopher Lee, UConn Health). The PRE and POST ABR waveforms were generated by averaging the responses to each ABR stimulus (1266 repetitions each) before and after the LDS and plotted relative to the onset of the ABR stimulus. ABRs were filtered between 500-3000 Hz. The peak and trough of each ABR wave were manually selected by a researcher blind to tinnitus status. Peak V was identified as the peak before the deepest trough in the ABR waveform. Other waves were identified by counting peaks in relation to their timing to wave V. In most recordings, wave VI could be identified and was included in analysis to quantify the entire sound-driven response. Wave amplitude was calculated by the absolute distance between the peak and the following trough. The peak latency relative to the onset of the ABR stimulus determined wave latency. Changes in ABR wave amplitude PRE and POST LDS were classified as potentiated or suppressed based on the results of the normalization of the PRE and POST results (POST-PRE/POST+PRE) referred to here as delta (Δ).

An automated analysis correlated a 12 ms long response window (2 ms - 14 ms after the stimulus onset) of PRE and POST responses without manual peak picking. Using bootstrapping, half of the PRE recordings from the selected time window were randomly selected and averaged to generate an ABR waveform. Then, the other half of the PRE recordings from the selected time window were averaged to generate a second ABR waveform. The two waveforms were then correlated to generate a correlation coefficient (R-value) for those PRE ABR waveforms (PRE:PRE). This process was repeated 100 times using 100 different random samples to generate 100 PRE:PRE R-values. The same analysis was repeated using the recordings POST LDS (POST:POST). Finally, PRE LDS and POST LDS responses were correlated using the same method (PRE:POST, also resulting in 100 R-values). This analysis of PRE:PRE, POST:POST, and PRE:POST was performed for each ABR_LDS_ recording. The mean and standard deviation of the distribution of PRE:PRE, POST:POST, and PRE:POST-R-values were calculated.

#### Analysis of extracellular ICC recordings

The presence of a long-duration sound-induced afterdischarge, a sustained increase in spontaneous firing rate after the LDS (Ono et al. 2016), was determined by comparing the spontaneous activity before the LDS (PRE LDS) and after the LDS (POST LDS) in each channel that responded to that LDS. Afterdischarge responses were characterized as present/not present in each electrode channel based on the comparison of POST LDS spiking activity to the PRE LDS spiking activity. POST spiking activity was only considered to be an afterdischarge if the POST spontaneous spike rate of the channel exceeded the 95% confidence interval of the PRE spontaneous spike rate for three consecutive bins of 2.5 s. Only afterdischarge responses that started within 30 seconds of the LDS offset were included.

The response to tone pips were only considered valid responses if the response started after 3.4 ms. The activity evoked by the PRE LDS and POST LDS tone pips was analyzed in channels with a sound-driven response to the LDS <u>and</u> either the PRE or POST tone pips. The same methods used in Burghard et al. (2022) were employed to calculate the overall spike rate and peristimulus time histogram (PSTH). The overall spike rate PRE was calculated using the PSTH of the entire cycle after tone pip presentation (approx. 47 ms) across all six 10 second tone pip trains before the LDS. The overall spike rate POST was calculated using the PSTH of the entire cycle after the tone pip presentation of the first 10 second tone pip train (T1) after the LDS. The PSTH from T1 was selected because Burghard et al. (2022) showed that in naïve mice, sound-driven activity in T1 showed the largest difference compared to sound-driven activity PRE LDS. For comparison and plotting, the difference in spiking activity was normalized within each channel ((POST —PRE)/(PRE+POST)) and reported as ‘delta’. Changes in sound-evoked response before and after the LDS were classified as potentiated or suppressed if these changes were positive or negative, respectively, after normalization. For each analysis, only responsive channels to the specific stimulus were used. Statistical analysis was done for each side (contra- or ipsilateral to the sound exposed ear) separately via one-way ANOVA (factor: tinnitus status – tinnitus, no-tinnitus, control) followed by a Scheffe post-hoc test where appropriate.

### Statistical tests

2-sample proportions tests were used to analyze differences in the percentage of LDS channels in ICC recordings. One-way analysis of variance (ANOVAs) were used to analyze spike count and response duration in the ICC recordings. Two-way ANOVAs were used to analyze ABR_LDS_ responses, with tinnitus status and ear (sound exposed or unexposed ear) as factors.

### Software accessibility

Access to manuals and software is available upon request, provided users agree to share data while the program is under development.

## 3 Results

### Prevalence of afterdischarge activity in mice with tinnitus

We have previously described a long-duration sound-induced afterdischarge in the inferior colliculus multi-unit activity (Burghard et al. 2022). To determine if afterdischarges occur more frequently in tinnitus, non-tinnitus, or control mice, we examined extracellular multichannel recordings from sound-responsive channels in both ICCs of unilaterally sound-exposed mice and the right ICC of control mice. (Fig. 1). Afterdischarges were characterized as present/not present based on the POST LDS spiking activity compared to the PRE LDS baseline activity. Following a long-duration sound, non-tinnitus mice had a lower percentage of channels with an afterdischarge than the ICC contralateral to the sound-exposed ear in tinnitus mice and right ICC in control mice. In contrast, tinnitus mice were not significantly different than control (Fig. 1A. 2 sample proportions test, contralateral ICC: Tinnitus vs. non-tinnitus, z = 2.255, p = 0.024. Tinnitus vs. control, z = 0.197, p=0.844. Non-tinnitus vs. control, z = 2.829, p=0.005). In contrast, in the ICC ipsilateral to the sound-exposed ear, both the tinnitus and non-tinnitus mice had increased numbers of afterdischarge channels overall than the control (Fig. 1A. 2 sample proportions test, ipsilateral ICC: Tinnitus vs. control, z= -6.211, p = 5.26e-10. Non-tinnitus vs. control, z = -5.247, p= 1.54e-10. Non-tinnitus vs. tinnitus, z = -0.731, p = 0.465).

**Fig. 1:**
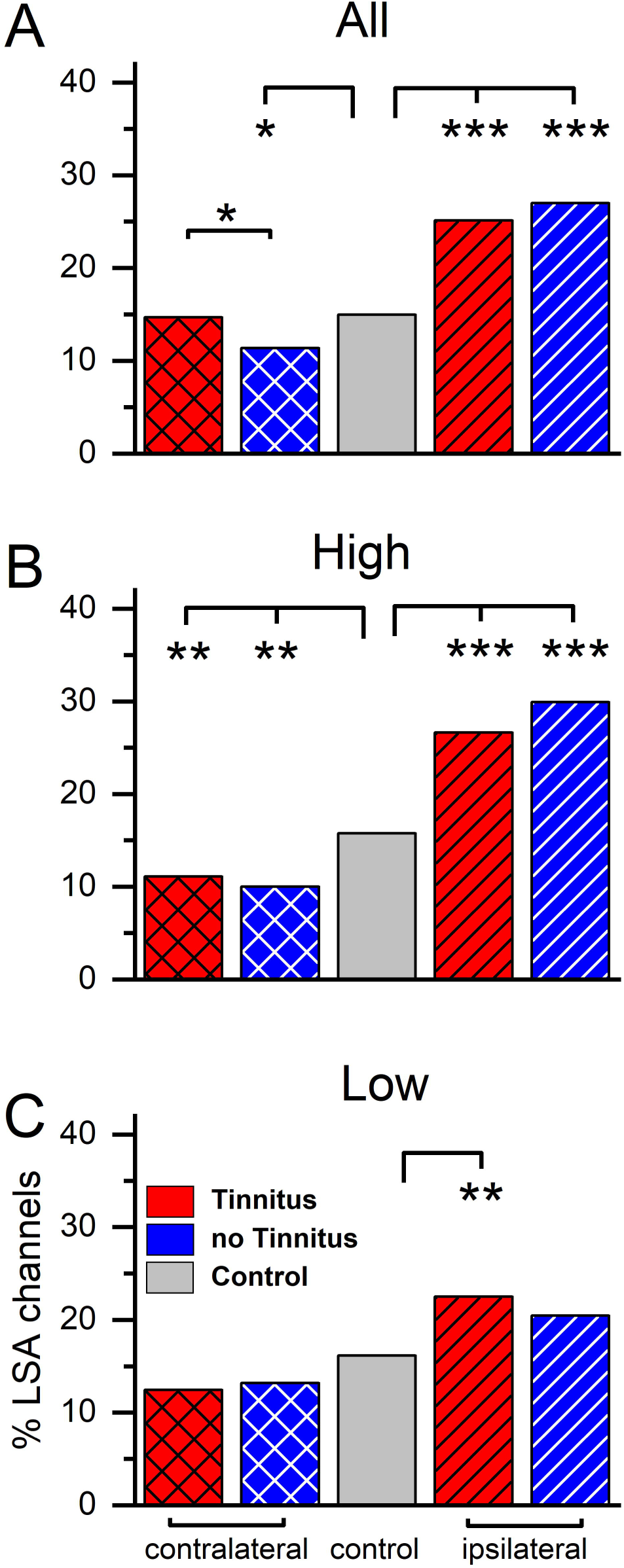
Sound exposure changes percentage of channels with long-duration sound-induced afterdischarge activity. Percent of afterdischarge (%LSA) positive channels separated by group and recording side. **A:** Percent afterdischarge positive channels when stimulated with an LDS (long-duration sound). Contralateral: tinnitus n=1055, non-tinnitus n=1062, control n=1999. Ipsilateral: tinnitus n=924, non-tinnitus n=426. **B:** Percent of afterdischarge positive channels using an LDS at or above 16 kHz (center of sound exposure frequency). Contralateral: tinnitus n=391, non-tinnitus n=309, control n=909. Ipsilateral: tinnitus n=365, non-tinnitus n=200. **C:** Percent of afterdischarge positive channels when using an LDS frequency below 16 kHz. Contralateral: tinnitus n=944, non-tinnitus n=753, control n=1054. Ipsilateral: tinnitus n=523, non-tinnitus n=219, control n=1054. Red: Tinnitus animals, blue: no tinnitus animals, grey: control animals. Cross hatch: recordings from the IC contralateral to the sound-exposed ear, stripes: recordings from the IC ipsilateral to the sound-exposed ear, no pattern: control mice, no sound exposure. *p<0.05, **p<0.01, ***p<0.001.

The unilateral sound exposure used to induce tinnitus was centered at 16 kHz. Based on previous studies (McFadden 1986, Cody and Johnstone 1981), more damage is expected in auditory regions tuned higher than the sound exposure center frequency than in the auditory areas tuned to lower frequencies. Therefore, we separated the trials by LDS center frequency, which was used to investigate if a frequency-specific effect was present.

Responses from high LDS center frequency stimuli (e.g. high, 16 kHz and above, Fig. 1B) resulted in a similar pattern to all stimuli combined. The percentage of afterdischarge channels in the contralateral ICC was not different between tinnitus, non-tinnitus, and control mice (2 sample proportions test: Tinnitus vs. control, z = 0.731, p=0.465. Non-tinnitus vs. control, z=-0.045, p=0.964. Non-tinnitus vs. tinnitus, z=0.628, p=0.529). On the ipsilateral ICC, sound-exposed animals showed significantly increased prevalence in afterdischarge channels compared to control animals (2 sample proportions test: Tinnitus vs. control, z=2.672, p=0.007. Non-tinnitus vs. control, z=4.491, p=7.077e-6. Tinnitus vs. non-tinnitus, z=-2.396, p=0.016).

Low LDS center frequency stimuli (e.g. low, <16 kHz, Fig. 1C) showed the smallest effect of sound exposure on both the contra- and the ipsilateral side. In the contralateral ICC, there was a significant effect of sound-exposure status, with tinnitus and non-tinnitus mice exhibiting a significantly smaller proportion of afterdischarge positive channels than control (2 sample proportion test: tinnitus vs. control, z=-4.857, p=1.19e-6. Non-tinnitus vs. control, z=-4.415, p=1.1e-6. Tinnitus vs. non-tinnitus, z=-0.189, p=0.849). Similarly, the trend of a higher afterdischarge prevalence after sound exposure was still present on the ipsilateral side for both tinnitus and non-tinnitus mice (2 sample proportions test: tinnitus vs. control, z=5.031, p=4.877e-7. Non-tinnitus vs. control, z=3.082, p=0.002. Tinnitus vs. non-tinnitus, z=0.454, p=0.649).

These data reflect a tinnitus-specific and a general plastic response in the ICC after unilateral sound exposure and hearing loss. In the ICC contralateral to the sound-exposed ear, tinnitus status affected the induction of afterdischarges. The effect was not frequency-specific but was general to all tested frequencies. This tinnitus-specific difference disappeared when the responses were separated by stimulus frequency. However, a frequency-specific effect of sound exposure was visible. High LDS center frequencies resulted in a more substantial sound exposure effect than low LDS center frequencies, supporting the notion that sound exposure predominately affects the sound exposure frequency and above.

### Effect of sound exposure and tinnitus status on stimulus-evoked responses in ICC

Because sound exposure and tinnitus status altered the percentage of afterdischarge channels after an LDS in the ICC, it follows that ICC activity in response to sounds may also be altered. To determine if sound exposure influenced sound-evoked responses, we quantified spike count and response duration of extracellular recordings to 3 ms tone pips before and after LDS. We first compared responses to tone pips <u>before the LDS</u> in sound-exposed and control mice (Fig. 2). Sound-exposed mice exhibited a significant increase in tone-driven spike count across all frequencies in both ICC compared to control mice (Fig. 2 A-C). Contra: One-way-ANOVA: F=64.58, p<0.0001.

**Fig. 2:**
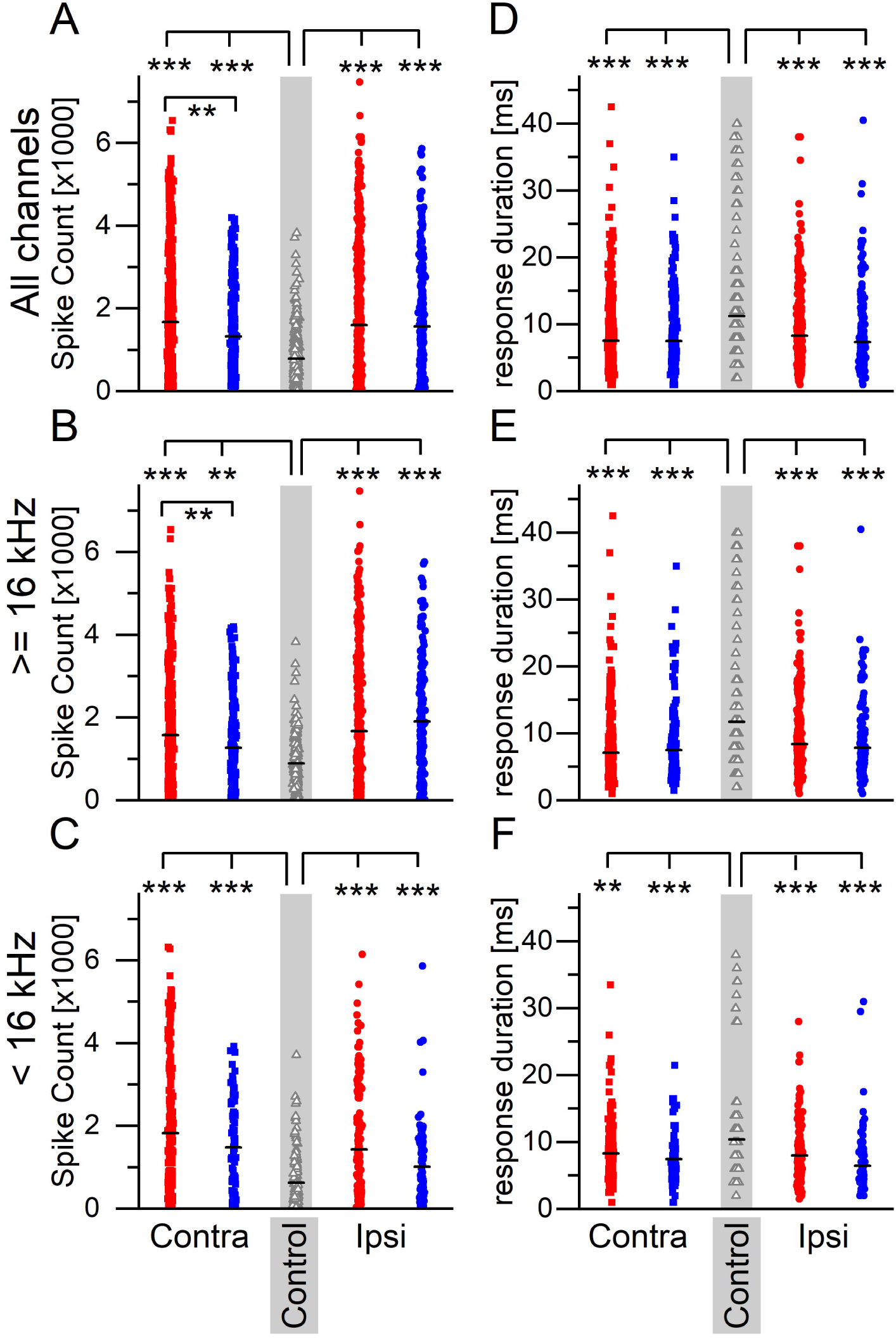
Sound exposure leads to an increase in firing activity in response to tone pips and a reduction in response duration compared to non-sound exposed controls. **A-C**: Spiking activity in response to 3 ms tone pips. **A)** Spiking activity to all tone pips. Contralateral: tinnitus n=635, non-tinnitus n=310, control n=360. Ipsilateral: tinnitus n=576, non-tinnitus n=285. **B:** Spiking activity in response to tone pips ≥ 16 kHz. Contralateral: tinnitus n=398, non-tinnitus n=227, control n=225. Ipsilateral: tinnitus n=402, non-tinnitus n=179. **C:** Spiking activity in response to tone pips < 16 kHz. Contralateral: tinnitus n=237, non-tinnitus n=83, control n=135. Ipsilateral: tinnitus n=174, non-tinnitus n=83. **D-F:** Response duration to same tone pips as in A-C. **D:** Contralateral: tinnitus n=635, non-tinnitus n=310, control n=360. Ipsilateral: tinnitus n=174, non-tinnitus n=106. **E:** Contralateral: tinnitus n=398, non-tinnitus n=227, control n=225. Ipsilateral: tinnitus n=576, non-tinnitus n=285. **F:** Contralateral: tinnitus n=237, non-tinnitus n=83, control n=135. Ipsilateral: tinnitus n=174, non-tinnitus n=106, control n=135. Red: Tinnitus animals, blue: no tinnitus animals, grey: control animals. Squares: recordings from the IC contralateral to the sound-exposed ear, circles: recordings from the IC ipsilateral to the sound-exposed ear, open triangles: control mice, no sound exposure. Gray background indicates recordings from control animals. *p<0.05, **p<0.01, ***p<0.001.

Scheffe post-hoc: tinnitus vs. non-tinnitus p=1.27e-4, tinnitus vs. control p<0.0001, non-tinnitus vs. control p=0.0001. Ipsi: One-way-ANOVA: F=48.561, p<0.0001. Scheffe post-hoc: tinnitus vs. non-tinnitus p=0.962, tinnitus vs. control p<0.0001, non-tinnitus vs. control p<0.0001). ‘High-frequency stimuli resulted in a tinnitus-specific increase in tone-driven spike count on the contralateral side only (Contra: One-way-ANOVA: F=28.255, p<0.0001. Scheffe post-hoc: tinnitus vs. non-tinnitus p=0.003, tinnitus vs. control p<0.0001, non-tinnitus vs. control p=0.001. Ipsi: One-way-ANOVA: F=33.502, p<0.0001. Scheffe post-hoc: tinnitus vs. non-tinnitus p=0.1632, tinnitus vs. control p<0.0001, non-tinnitus vs. control p<0.0001). There were no tinnitus-specific changes in spike count using low-frequency stimuli (Contra: One-way-ANOVA: F=37.688, p<0.0001. Scheffe post-hoc: tinnitus vs. non-tinnitus p=0.113, tinnitus vs. control p<0.0001, non-tinnitus vs. control p<0.0001. Ipsi: One-way-ANOVA: F=15.84, p<0.0001. Scheffe post-hoc: tinnitus vs. non-tinnitus p=0.085, tinnitus vs. control p=7.797e-4, non-tinnitus vs. control p<0.0001). Interestingly, the response duration was shorter in the unilaterally sound-exposed animals than in control animals, also regardless of the recording side (Fig. 2D-F. All frequencies; Contra: One-way-ANOVA: F=53.554, p<0.0001. Scheffe post-hoc: tinnitus vs. non-tinnitus p=0.983, tinnitus vs. control p<0.0001, non-tinnitus vs. control p<0.0001. Ipsi: One-way-ANOVA: F=40.313, p<0.0001. Scheffe post-hoc: tinnitus vs. non-tinnitus p=0.101, tinnitus vs. control p<0.0001, non-tinnitus vs. control p<0.0001. High frequencies; Contra: One-way-ANOVA: F=46.362, p<0.0001. Scheffe post-hoc: tinnitus vs. non-tinnitus p=0.749, tinnitus vs. control p<0.0001, non-tinnitus vs. control p<0.0001. Ipsi: one-way-ANOVA: F=26.575, p<0.0001. Scheffe post-hoc: tinnitus vs. non-tinnitus p=0.632, tinnitus vs. control p<0.0001, non-tinnitus vs. control p<0.0001. Low frequencies; Contra: One-way-ANOVA: F=10.028, p<0.0001. Scheffe post-hoc: tinnitus vs. non-tinnitus p=0.449, tinnitus vs. control p=0.001, non-tinnitus vs. control p=3.58e-4. Ipsi: One-way-ANOVA: F=15.84, p<0.0001. Scheffe post-hoc: tinnitus vs. non-tinnitus p=0.085, tinnitus vs. control p<7.797e-4, non-tinnitus vs. control p<0.0001).

These data suggest that unilateral sound exposure generally increases sound-driven spike count and decreases response duration in both ICCs. At the same time, tinnitus coincides with further spike rate increases only in the ICC contralateral to the sound exposure.

### Effect of sound exposure and tinnitus status on LDS-driven plasticity

To determine if there was a tinnitus-specific LDS effect on sound-evoked plasticity, the change in tone-evoked spike count and response duration before (PRE) and after (POST) the LDS were compared (Fig. 3). Mean delta (normalized PRE:POST difference) spike counts below zero indicate that the sound-driven spiking activity was suppressed after the LDS. In the ICC contralateral to the sound exposed ear, non-tinnitus mice had a significantly lower delta (normalized PRE:POST difference) spike count than tinnitus and control mice for all frequencies, including when separating responses to high and low-frequency stimuli (Fig. 3 A-C. All frequencies: One-way-ANOVA F=12.149, p<0.0001. Post-hoc Scheffe, tinnitus vs. non-tinnitus p=0.0001, tinnitus vs. control p=0.957, non-tinnitus vs. control p=1.167e-4. High frequencies: One-way-ANOVA F=7.124, p=8.60e-4. Post-hoc Scheffe, tinnitus vs. non-tinnitus p=0.003, tinnitus vs. control p=0.951, non-tinnitus vs. control p=0.005. Low frequencies: One-way-ANOVA F=5.099, p=0.006. Post-hoc Scheffe, tinnitus vs. non-tinnitus p=0.011, tinnitus vs. control p=0.998, non-tinnitus vs. control p=0.022). These data suggest that sound-driven activity in non-tinnitus mice is more suppressed after the LDS than in tinnitus and control mice. Interestingly, there was no significant difference in delta spike count between tinnitus and control. In the ICC ipsilateral to the sound exposed ear, a tinnitus-specific effect on delta spike count was also present when looking at all frequencies and responses to high-frequency stimuli but not low-frequency stimuli (Fig. 3A. All frequencies: One-way-ANOVA F=8.459, p=2.26e-4. Post-hoc Scheffe, tinnitus vs. non-tinnitus p=0.037, tinnitus vs. control p=0.118, non-tinnitus vs. control p=2.27e-4. High frequencies: One-way-ANOVA F=3.586, p=0.028. Post-hoc Scheffe, tinnitus vs. non-tinnitus p=0.751, tinnitus vs. control p=0.028, non-tinnitus vs. control p=0.312. Low frequencies: One-way-ANOVA F=10.887, p<0.0001. Post-hoc Scheffe, tinnitus vs. non-tinnitus p=1.88e-4, tinnitus vs. control p=0.998, non-tinnitus vs. control p=3.07e-4).

**Fig. 3:**
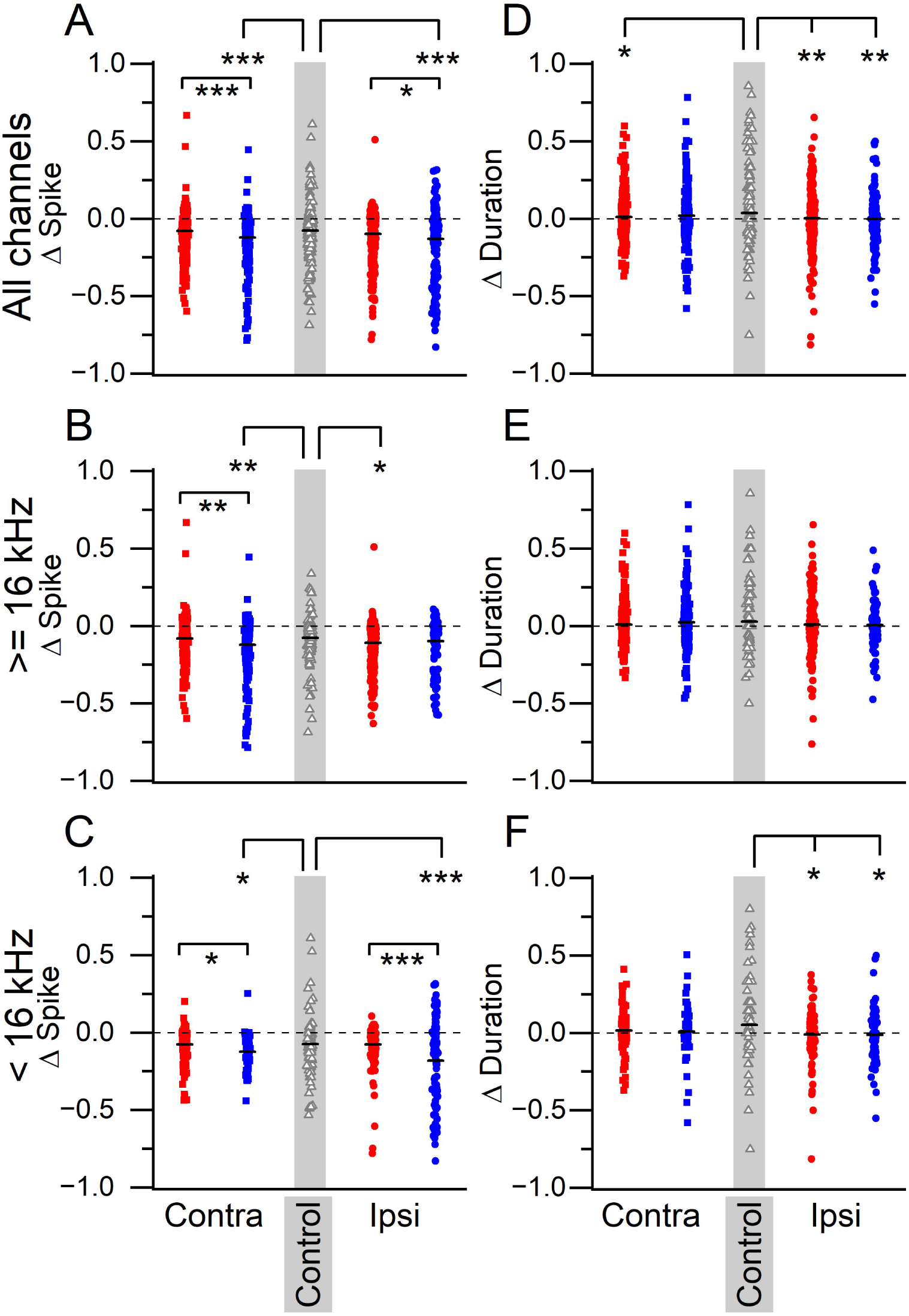
LDS-induced plasticity reveals tinnitus status-dependent differences. **A-C**: Normalized (delta) difference in spiking activity in response to 3 ms tone pips before and after a long-duration sound. **A:** Spiking activity to all tone pips. Contralateral: tinnitus n=573, non-tinnitus n=284, control n=319. Ipsilateral: tinnitus n=500, non-tinnitus n=254**. B:** Spiking activity in response to tone pips ≥ 16 kHz. Contralateral: tinnitus n=359, non-tinnitus n=205, control n=202. Ipsilateral: tinnitus n=357, non-tinnitus n=156. **C:** Spiking activity in response to tone pips < 16 kHz. Contralateral: tinnitus n=214, non-tinnitus n=79, control n=117. Ipsilateral: tinnitus n=143, non-tinnitus n=98. **D-F:** Normalized difference in response duration to same tone pips as in A-C. **D:** Contralateral: tinnitus n=573, non-tinnitus n=284, control n=319. Ipsilateral: tinnitus n=500, non-tinnitus n=254. **E:** Contralateral: tinnitus n=359, non-tinnitus n=205, control n=202. Ipsilateral: tinnitus n=357, non-tinnitus n=205. **F:** Contralateral: tinnitus n=214, non-tinnitus n=79, control n=117. Ipsilateral: tinnitus n=143, non-tinnitus n=98. Δ = normalized difference ((POST-PRE)/(POST+PRE)). Red: Tinnitus animals, blue: no tinnitus animals, grey: control animals. squares: recordings from the IC contralateral to the sound-exposed ear, circles: recordings from the IC ipsilateral to the sound-exposed ear, open triangles: control mice, no sound exposure. Gray background indicates recordings from control animals. *p<0.05, **p<0.01, ***p<0.001.

We also wanted to determine if there was an effect of tinnitus status on delta duration (Fig. 3 D-E). When looking at responses across all frequencies (Fig. 3 D), in the contralateral ICC, tinnitus mice had similar delta response durations to control mice, exhibiting a tinnitus effect (All frequencies; One-way-ANOVA F=3.259, p=0.039. Post-hoc Scheffe, tinnitus vs. non-tinnitus p=0.697, tinnitus vs. control p=0.038, non-tinnitus vs. control p=0.359.) In contrast, tinnitus status did not affect delta duration in the contralateral IC when the responses to high or low frequency stimuli were separated (High frequencies: One-way-ANOVA F=1.39, p=0.249. Low frequencies: One-way ANOVA F=2.992, p=0.051). Showing that sound exposure, but not tinnitus status, affected the change in tone-evoked response duration in the ipsilateral ICC (Fig. 3D) if stimulus frequency was not a factor. On the ipsilateral side, both sound-exposed groups tended to show more suppression of duration in comparison to control animals (Fig. 3 D-E. All frequencies: One-way-ANOVA F=6.498, p=0.002. Post-hoc Scheffe, tinnitus vs. non-tinnitus p=0.939, tinnitus vs. control p=0.006, non-tinnitus vs. control p=0.009. High frequencies: one-way-ANOVA F=1.641, p=0.194. Low frequencies: One-way-ANOVA F=5.81, p=0.003. Post-hoc Scheffe, tinnitus vs. non-tinnitus p=0.995, tinnitus vs. control p=0.01, non-tinnitus vs. control p=0.017).

### Effect of tinnitus on ABR latency and amplitude

To determine if changes in the baseline ABR were associated with tinnitus, we compared the amplitude and latency of tone-evoked ABR wave I and wave V from tinnitus, non-tinnitus, and control mice (Fig. 4). The average wave I amplitude of the tone-evoked PRE-LDS waveforms from the ABR_LDS_ were measured from both the sound-exposed and un-exposed (i.e., ear-plugged during sound exposure) ears from tinnitus mice (TE and TU, respectively), sound-exposed and un-exposed ears from non-tinnitus mice (NTE and NTU, respectively), and control mice (C, Fig. 4A). Control mice had a significantly larger wave I amplitude compared to all other cohorts, except for NTU ears (Fig. 4A. 2-way ANOVA F(5,137) = 3.96, p=0.002. Post-hoc Scheffe test: C vs. TE p=0.002, C vs. TU p=0.02, C vs. NTE p=0.003, C vs. NTU p=0.079). These results showed that sound exposure significantly reduced the amplitude of ABR wave I. The eighth cranial nerve is considered the wave I generator (Eggermont 2019), so sound exposure may have caused cochlear or spiral ganglion neuron damage that reduced wave I amplitude. There were no significant differences in ABR wave V amplitude between any category (Fig. 4B. 2-way ANOVA F(5,138) = 0.63, p=0.670). This finding is consistent with the notion of compensation in higher auditory brainstem regions for sound-exposure damage in the periphery.

**Figure 4:**
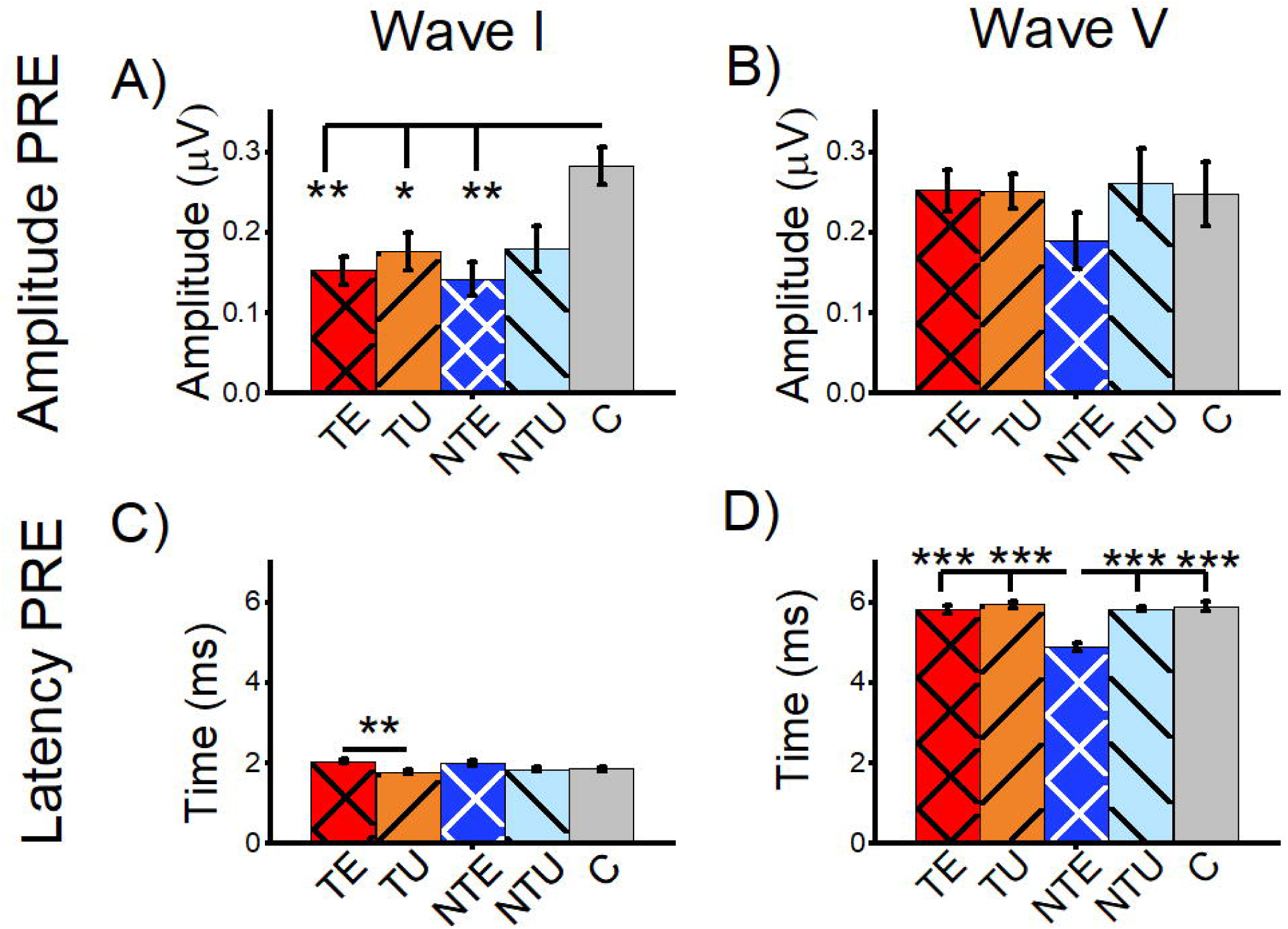
Sound exposure affected amplitude of wave I and latency of wave V of tone-evoked ABRs before the long duration sound. All data are mean and standard error of the mean. Data are from tone-evoked ABRs before the long-duration sound (PRE LDS). **A:** Average amplitude for tone-evoked wave I. **B:** Average amplitude for tone-evoked wave V. **C:** Average latency of tone-evoked wave I. **D:** Average latency of tone-evoked wave V. TE = tinnitus exposed ears (red, crosshatch). TU = tinnitus unexposed ears (orange, black stripes). NTE = non-tinnitus exposed ears (blue, white crosshatch). NTU = non-tinnitus unexposed ears (light blue, black stripes). C = control, unexposed (grey bars). * = p<0.05, ** = p<0.01, *** = p<0.001. TE n=40, TU n=41, NTE n=22, NTU n= 21, C n=21

In tinnitus mice, a small but significant increase in wave I latency in the sound-exposed ear was observed compared to the un-exposed ear, with no significant differences in other cohorts (Fig. 4C. 2-way ANOVA F(5,137) = 3.91, p=0.002. Post-hoc Scheffe test: TE vs. TU p=0.002). In contrast, sound-exposed non-tinnitus mice exhibited a significantly shorter wave V latency compared to all other cohorts, with no other significant differences seen (Fig. 4D. 2-way ANOVA F(5,137) = 10.64, p=0.0004. Post-hoc Scheffe test: NTE vs. TE p=5.4e-8, NTE vs. TU p=2.2e-8, NTE vs. NTU p=7.9e-7, NTE vs. C p=1.5e-7).

### Correlation analysis of the ABR_LDS_

Traditional methods of ABR analysis rely on manual peak picking that may be subjective (Eggermont 2019). Automated ABR analysis may be more objective. Therefore, we implemented an automated analysis method for the ABR_LDS_ and compared ABR waveforms before and after the LDS using a bootstrapping correlation analysis (Fig. 5). An example of a bootstrapping correlation analysis readout is shown in Figure 7A. Each trace representing an averaged PRE-LDS sample is shown in blue, and the POST-LDS traces are red (Fig. 5A, top panel). In this image, the PRE- and POST-LSD waveforms differ in shape. The bottom panel (Fig. 5A) shows the distribution of the correlation coefficients (R-values) from each PRE:PRE (Fig. 5A, bottom panel, blue), POST:POST (Fig. 5A, bottom panel, red), and PRE:POST (Fig. 5A, bottom panel, green) of the traces in the top panel. In this example, the PRE:PRE and POST:POST mean R-values were high, suggesting that the PRE and POST ABRs were internally consistent. In contrast, the mean R-value PRE:POST distribution was reduced, indicating that the PRE and POST ABR_LDS_ waveforms were markedly different.

**Figure 5:**
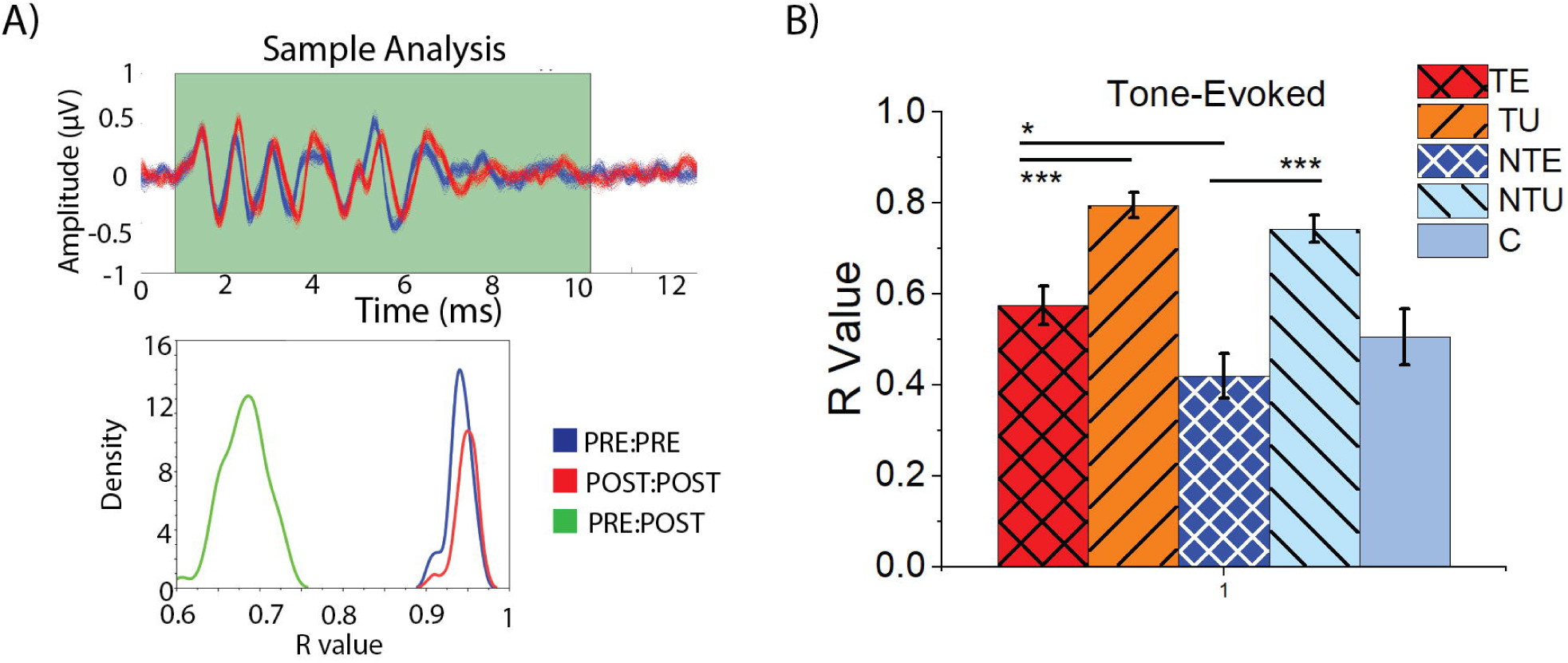
ABRs from non-tinnitus sound exposed ears were more affected by the LDS than tinnitus ears using bootstrapping correlation analysis. **A:** Example correlation analysis of one ABR-NSP response. Top: plot of bootstrapped ABR waveforms, blue represents PRE data, red represents POST data. Bottom: Graph showing the distribution of PRE:PRE (blue), POST:POST (red), and PRE:POST (green) R-values. **B:** Mean and standard error of the mean of PRE:POST R-value for tone-evoked ABRs. TE = tinnitus exposed ears (red, crosshatch). TU = tinnitus unexposed ears (orange, black stripes). NTE = non-tinnitus exposed ears (blue, white crosshatch). NTU = non-tinnitus unexposed ears (light blue, black stripes). C = control, unexposed (grey bars). * = p<0.05, ** = p<0.01, *** = p<0.001. R-Value = correlation coefficient. p = p-value. PRE = before the LDS, POST = after the LDS. ABR = auditory brainstem response, LDS = long-duration sound. TE n=38, TU n=37, NTE n=20, NTY n=23, C n=16.

To determine if the LDS affected the tone-evoked ABR_LDS_ waveforms in a tinnitus status-dependent manner, the mean R-value PRE:POST was compared across all cohorts (Fig. 5B). In both the tinnitus and non-tinnitus mice, the average responses driven by the sound-exposed ear exhibited significantly lower mean R-values compared to the un-exposed ear, suggesting that sound exposure reduced the PRE:POST ABR_LDS_ correlation (Fig. 5B. 2-way ANOVA: F(5,129) = 19.16, p=2.2e-12. Post-hoc Scheffe, TE vs. NTE p=0.046, NTE vs. NTU p=3.9e-8). Furthermore, responses driven by the tinnitus sound-exposed ear exhibited a significantly larger mean R-value than non-tinnitus sound-exposed ears, suggesting that tinnitus may compensate for sound exposure-related deficits (Post-hoc Scheffe, TE vs. TU p=6.9e-5). However, none of these responses were significantly different from control.

### Comparison of ABR waveform metrics in the ABR_LDS_

Tinnitus has been linked to increased central gain, defined as the compensatory increase in neural activity in the central auditory system in response to the loss of peripheral input (Auerbach et al. 2014), Amplitude ratios between peripheral and central ABR waves have been used to measure central gain in animal models (Cai et al. 2018, Parthasarathy and Kujawa 2018, Mohrle et al. 2016). Therefore, we calculated the wave V-I and V-III amplitude ratios for both PRE-LDS and POST-LDS ABR_LDS_ waveforms and found no significant differences between any of the tested groups (PRE-LDS: V/I ratio: 2-way ANOVA, F(5,126) = 1.23, p=0.099. V/III ratio: 2-way ANOVA, F(5, 128) = 1.21, p=0.307. POST-LDS: V/I ratio: 2-way ANOVA, F(5,124) = 2.06, p=0.074. V/III ratio: 2-way ANOVA, F(5, 122) = 1.99, p=0.084).

Increased ABR inter-wave latency has been reported in human patients with tinnitus (Singh et al. 2011, Kehrle et al. 2008). Therefore, we calculated the inter-wave latency between waves I and V (I-V) or VI (I-VI) for both PRE-LDS and POST-LDS ABR_LDS_ waveforms, again finding no significant difference between tinnitus, non-tinnitus, and control mice. (I-V latency: PRE; 2-way ANOVA, F(5,123) = 1.94, p=0.091. POST; 2-way ANOVA, F(5,123) = 1.57, p=0.173. I-VI latency: PRE; 2-way ANOVA, F(5,118) = 2.27, p=0.0513. POST; 2-way ANOVA, F(5,118) = 2.18, p=0.059).

ABR_LDS_ correlation analysis showed that the LDS resulted in tinnitus-specific changes in the ABR waveform. Therefore, we also analyzed the difference in PRE-LDS and POST-LDS ABR amplitude ratios and inter-wave latencies to determine the effect of the LDS (Fig 6). There were not LDS-induced changes on either amplitude ratios (V/I Difference: 2-way ANOVA, F(5,125) = 1.71, p=0.137. V/III Difference: 2-way ANOVA, F(5, 127) = 1.69, p=0.140) or inter-wave latencies (I-V Difference; 2-way ANOVA, F(5,123) = 2.03, p=0.078. I-VI latency: Difference; 2-way ANOVA, F(5,118) = 0.50, p=0.773). Therefore, the tinnitus-specific changes induced by the LDS were not evident in central gain and inter-wave latency measures.

**Figure 6:**
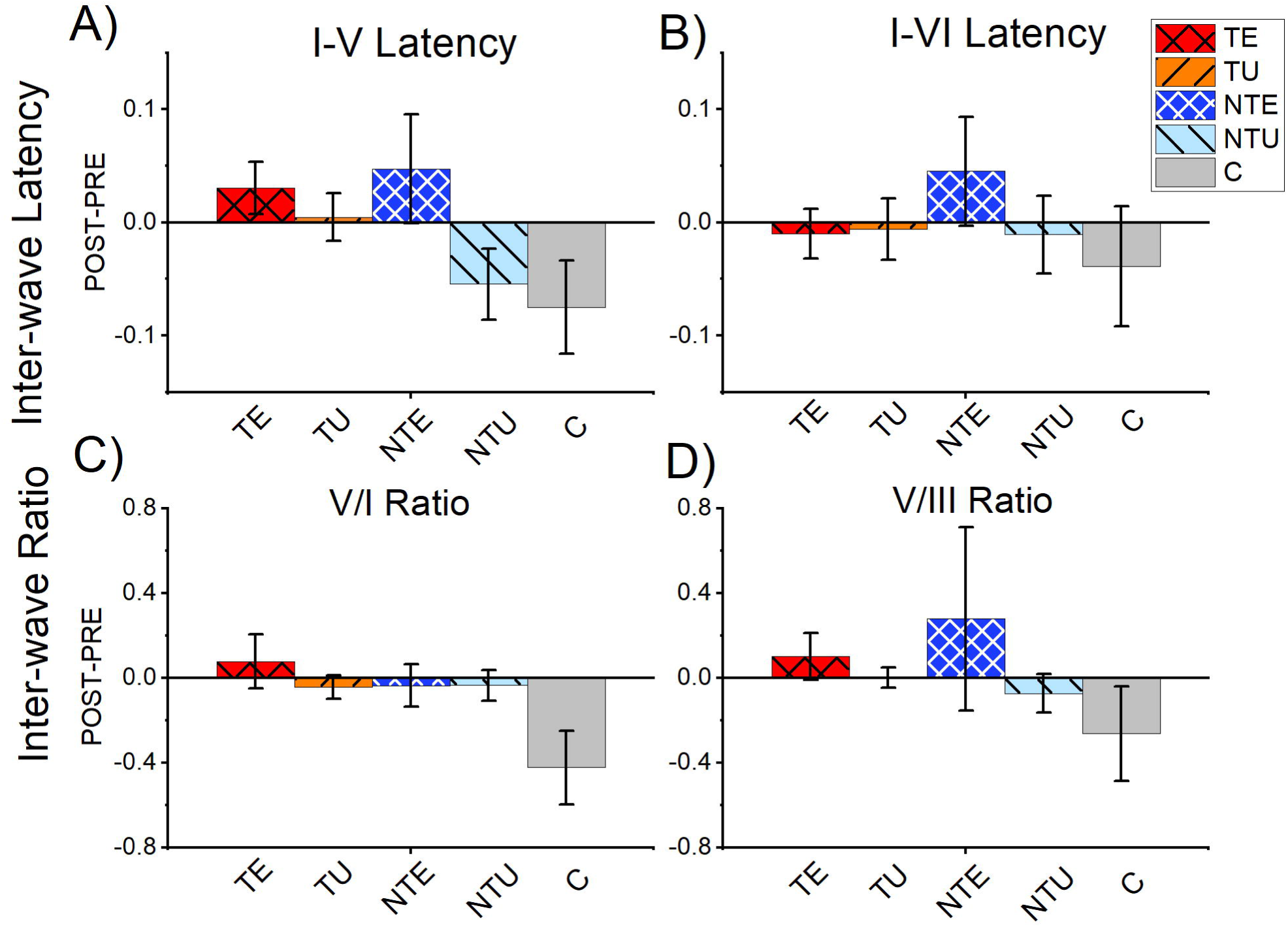
There were no significant differences in amplitude ratio or inter-wave latency from tone-evoked ABRs. All data are mean and standard error of the mean from tone-evoked ABRs. **A:** Difference of PRE LDS I-V latency subtracted from POST LDS I-V latency. **B:** Difference of PRE LDS I-VI latency subtracted from POST LDS I-VI latency. **C)** Difference of the PRE LDS V:I ratio subtracted from the POST LDS V:I ratio. **D)** Difference of the PRE LDS V:III ratio subtracted from the POST LDS V:III ratio. TE = tinnitus exposed ears (red, crosshatch). TU = tinnitus unexposed ears (orange, black stripes). NTE = non-tinnitus exposed ears (blue, white crosshatch). NTU = non-tinnitus unexposed ears (light blue, black stripes). C = control, unexposed (grey bars). * = p<0.05, ** = p<0.01, *** = p<0.001. ABR = auditory brainstem response. TE n=39, TU n=39, NTE =22, NTU n=21, C n=14.

### Differences in peak amplitudes in the ABR_LDS_

The bootstrapping correlation analysis revealed a difference in the PRE and POST ABR_LDS_ waveforms. However, manual peak-picking analysis was needed to determine the affected waves. As outlined in the methods, the delta peak-peak amplitude was calculated for the tone-evoked ABR_LDS._

A sample ABR_LDS_ waveform taken before the LDS (PRE, blue) and after the LDS (POST, red) shows the LDS’s influence on the ABR_LDS_ peaks (Fig. 7A). Figure 8B shows the average delta amplitude for all ABR_LDS_ waves from all tested frequencies and all cohorts. Consistent with results from the bootstrapping correlation analysis, the tone-evoked ABRs from sound-exposed ears in tinnitus and non-tinnitus mice were significantly different, with a significant suppression evident in non-tinnitus mice (Fig. 7B. 2-way ANOVA: F(5, 831) = 3.8, p=0.049. Post-hoc Scheffe test, p=0.047). Notably, the delta ABR_LDS_ amplitude difference in control mice was intermediate to the data from both cohorts of sound-exposed mice. It was not significantly different from either cohort. There were no significant differences in the delta ABR_LDS_ amplitude in responses from the unexposed ears for tinnitus and non-tinnitus mice.

**Figure 7:**
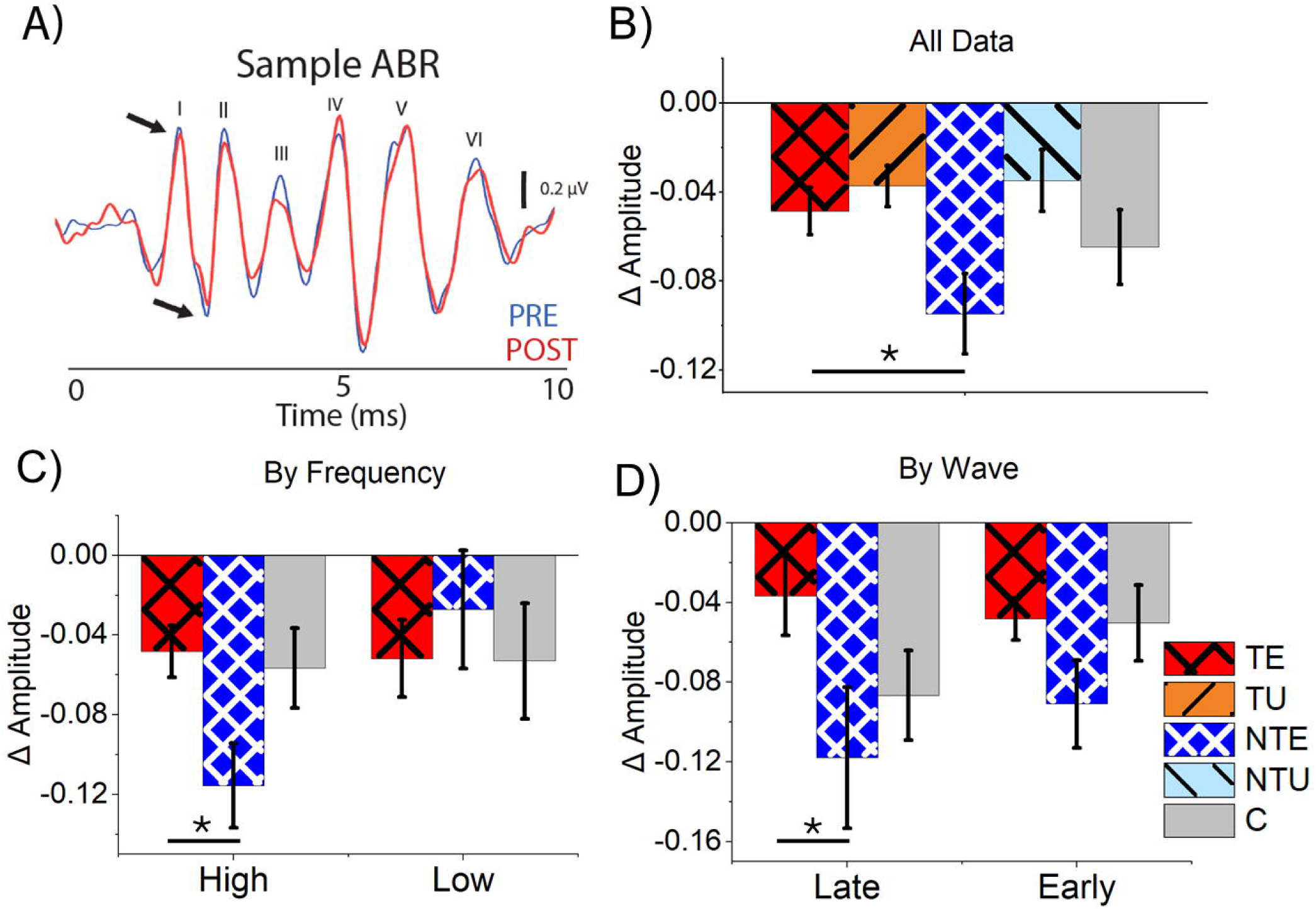
Tone-evoked ABRs at late waves and high stimulus frequencies were more suppressed in non-tinnitus mice compared to tinnitus mice. **A:** Example tone-evoked ABR waveform PRE and POST LDS with visualized waves labeled. Blue is PRE ABR data; red is POST ABR data. Black arrows indicate the peak and trough of wave I. **B:-D:** Data are mean and standard error of the mean. **B:** Normalized (delta) ABR amplitude difference for tone-evoked ABR_LDS_. TE n=250, TU n=264, NTE n=118, NTU n=123. **C)** Normalized ABR amplitude for tone-evoked ABRs for high (=>16 kHz) and low (<16 kHz) frequencies. High: TE n=170, NTE n=76, C n=60. Low: TE n=76, NTE n=47, C n=38. **D:** Normalized ABR amplitude for tone-evoked ABRs for late (IV, V, VI) and early (I, II, III) ABR waves. Late: TE n=87, NTE n=37, C n=30. Early: TE n=270, NTE n=60, C n=44. TE = tinnitus exposed ears (red, crosshatch). TU = tinnitus unexposed ears (orange, black stripes). NTE = non-tinnitus exposed ears (blue, white crosshatch). NTU = non-tinnitus unexposed ears (light blue, black stripes). C = control, unexposed (grey bars). * = p<0.05, ** = p<0.01, *** = p<0.001. LDS = long duration sound, NBN = narrow-band noise, PRE = before the LDS, POST = after the LDS, ABR = auditory brainstem response. Low = ABR and LDS frequencies below 16 kHz, high = ABR and LDS frequencies at or above 16 kHz, late = ABR waves IV, V, and VI, and early = ABR waves I, II, III. Δ = normalized differences (POST-PRE/POST+PRE).

**Figure 8:**
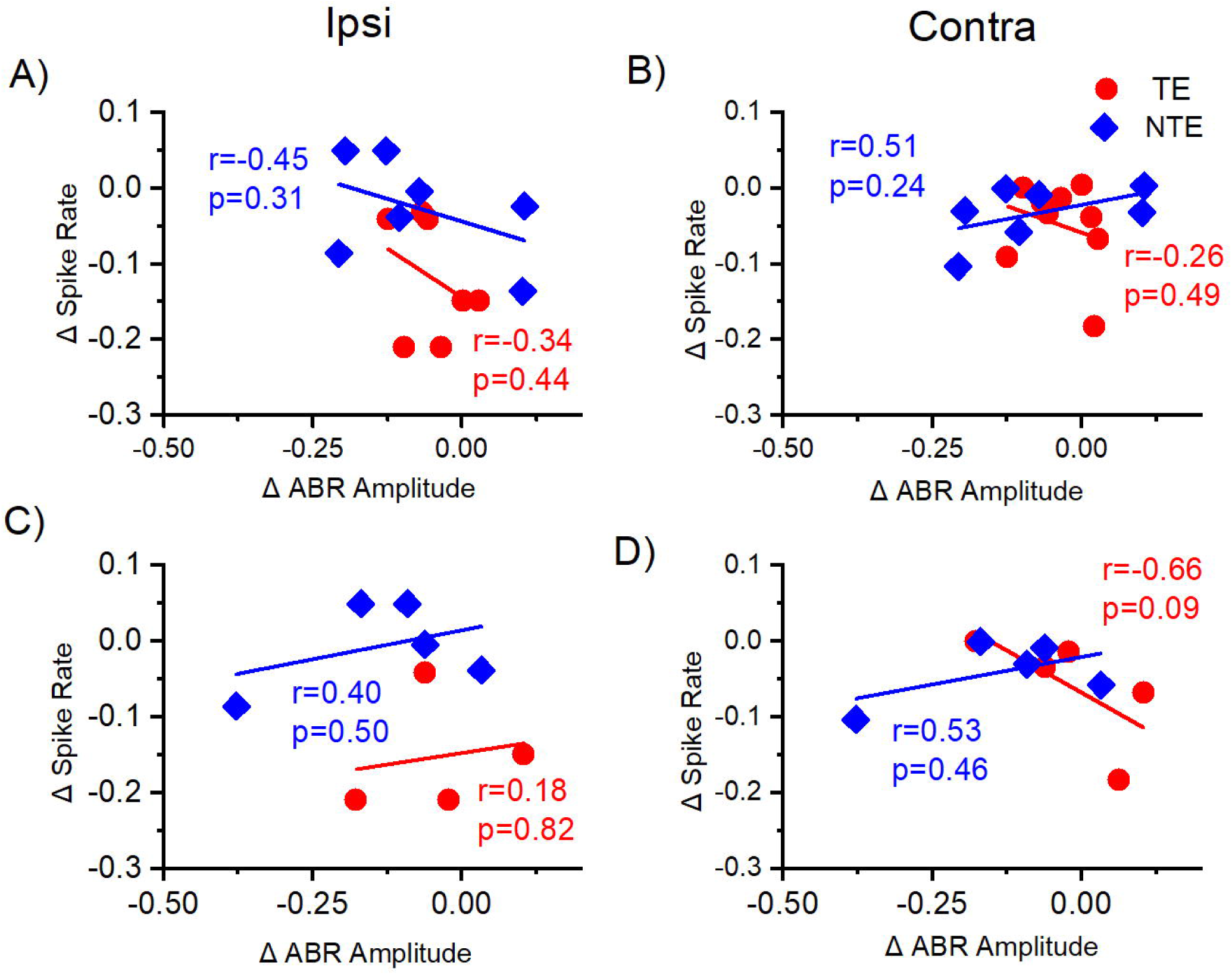
Correlation of normalized sound-driven spike rate from the ICC and ABR_LDS_ amplitude difference with tone stimuli. Correlation of normalized (delta) tone-driven spike rate recorded from the central nucleus of the inferior colliculus (ICC) (y-axis) to the normalized (delta) of tone-driven ABR amplitude from the exposed ear (x-axis) from same mouse with auditory stimulus at the same frequency. Each data point represents one auditory center frequency from one mouse. **A-B:** Correlation of the delta (Δ) tone-driven spike rate from the ICC to the average delta amplitude across all ABR waves from the exposed ear with all auditory stimulus frequencies**. A:** Ipsilateral ICC data correlated with the ABR amplitude difference from the exposed ear. **B:** Contralateral ICC data correlated with the ABR amplitude difference from the exposed ear. **C-D:** Correlation of the delta (Δ) spike rate from the ICC to the delta amplitude difference from late ABR waves (IV, V, VI) from the exposed ear with high-frequency auditory stimuli. **C:** Ipsilateral ICC data correlated with the ABR amplitude difference from the exposed ear. **D:** Contralateral ICC data correlated with the ABR amplitude difference from the exposed ear. TE = tinnitus (red, circle), NTE = non-tinnitus (blue, diamond). ABR = auditory brainstem response, r = Pearson’s correlation coefficient, p = p-value, Ipsi = ipsilateral ICC to the sound-exposed ear, Contra = contralateral ICC to the sound-exposed ear. Δ = normalized differences (POST-PRE/POST+PRE). TE n=9, NTE n=7.

Because we found a tinnitus-specific effect of stimulus frequency on sound-evoked plasticity in the ICC, the ABR_LDS_ results were separated by high (≥16 kHz) and low (<16 kHz) stimulus frequencies. At low ABR_LDS_ frequencies, there was no significant difference in average delta ABR amplitude between the cohorts (Fig. 7C. 1-way ANOVA: F(2,151) =1.57, p=0.167). However, the average delta amplitude from non-tinnitus sound-exposed ears was significantly suppressed compared to tinnitus sound-exposed ears in response to high-frequency stimuli (Fig. 7C. 1-way ANOVA, F(2,305) = 4.25, p=0.015. Post-hoc Scheffe test, TE vs. NTE p=0.017). Despite a tinnitus status-dependent effect, no significant differences existed between any sound-exposed cohort and control mice at both high and low stimulus frequencies. These results suggest that tinnitus status has a frequency-specific effect on sound-evoked plasticity.

Because the LDS affects sound-driven responses in the ICC, the data from early (I, II, III) and late (IV, V, VI) ABR_LDS_ waves were separately analyzed and compared to determine if the LDS had a wave-specific effect. The amplitude of early waves showed no significant differences between cohorts (Fig. 7D. 1-way ANOVA: F(2, 349) = 0.79, p-0.616). In contrast, the average late wave delta amplitude was significantly decreased in the non-tinnitus SE ears compared to the tinnitus SE ears (Fig. 7D. 1-way ANOVA: F(2,151) = 3.43, p=0.035. Post-hoc Scheffe test, TE vs. NTE p=0.047). Because the auditory midbrain drives later ABR_LDS_ waves, these results support our hypothesis that the LDS primarily affects sound-driven responses in the auditory midbrain and has a tinnitus-specific effect.

### Correlation of extracellular IC recordings and evoked ABRs

A subpopulation of mice underwent both extracellular ICC and ABR recordings to the LDS test (four tinnitus mice and three non-tinnitus mice), so it was possible to directly correlate the delta sound-driven spike rate in the contralateral and ipsilateral ICC with the average delta ABR amplitude from the sound-exposed ear (Fig. 8). It is worth noting that different anesthetics (ketamine for ABRs and isoflurane for extracellular ICC recordings) were used for each recording that may affect this comparison. Delta ABR_LDS_ amplitude was averaged across all waves. Figure 8A-B shows the correlation between the delta ABR_LDS_ amplitude from the exposed ear and the delta LDS test sound-driven spiking activity from left and right ICC, which were ipsilateral and contralateral to the sound-exposed ear, respectively. Although neither correlation is significant, the direction of the correlation is altered between the ipsilateral (negative correlation) and contralateral (positive correlation) ICC in non-tinnitus animals. In contrast, the tinnitus mice had a negative correlation in both ICCs. This was a surprising result because we expected that amplitude differences in the ABR_LDS_ would directly reflect ICC activity and that the results from these two electrophysiological recordings would be highly correlated.

It is a possibility that correlations were masked because of other auditory brainstem regions reflected in the ABR_LDS_. Therefore, we limited our comparison to the delta LDS test sound-driven spiking activity in the ICC and the late wave delta ABR_LDS_ amplitude from high-frequency stimuli (Fig. 8C-D). We chose to compare the data this way because the high-frequency stimuli data revealed tinnitus-specific differences and the late waves correspond to ICC activity. However, none of the correlations were significant. Both ipsilaterally and contralaterally, the non-tinnitus mice exhibited a positive correlation between the two electrophysiological recordings, while tinnitus mice only had a positive correlation in the ipsilateral ICC. These results are again surprising and suggest that auditory responses other than those from the ICC affect the ABR_LDS._

## 4 Discussion

This study aimed to reveal tinnitus-specific differences in sound-evoked neuronal activity before and after the LDS, measured with either ICC extracellular recordings or ABRs using the LDS test method. In the contralateral ICC, the tinnitus mice exhibited a significant increase in afterdischarge activity and tone-driven spike count compared to non-tinnitus mice. A comparison of the tone-driven spike rate before and after the LDS in the ICC revealed that sound-driven activity in non-tinnitus mice was more suppressed after the LDS than in tinnitus and control mice. Furthermore, the ABR_LDS_ revealed tinnitus-specific differences in delta late wave amplitude using high-frequency stimuli. These results suggest that comparing ABR_LDS_ waveforms before and after the LDS can reveal tinnitus-specific differences in mice after noise-induced hearing loss and may be a promising non-invasive electrophysiological method for identifying tinnitus. We theorize that sound exposure may result in suppression after the LDS but that tinnitus may ‘rescue’ sound-evoked responses.

### Using AA to identify behavioral evidence of tinnitus

We selected AA, an operant conditioning test, to identify mice with and without behavioral evidence of tinnitus. We have previously published evidence that tinnitus in AA is associated with increased spontaneous activity in the IC, a neural correlate of tinnitus, while a gap-detection-based tinnitus assessment is not (Fabrizio-Stover et al. 2022). The duration of the stimulus presentation in AA (5 seconds before shock) ensures that temporal processing deficits would not be reflected in the AA result. Testing over multiple weeks also ensures that any tinnitus behavior is sustained. The AA results are not produced by a bilateral frequency-specific hearing loss since, in these experiments, the mice have only unilateral hearing loss and normal thresholds in one ear. Therefore, although it is impossible to say with any certainty that a specific tinnitus assessment in laboratory animals is foolproof, we believe that AA is more likely to be a valid tinnitus assessment in mice than other methods.

Differences between extracellular IC LDS recordings and ABR_LDS_ recordings We found no significant correlations between ICC extracellular recordings and ABR_LDS_ recordings in the LDS-induced changes in sound-evoked activity. This finding is surprising, as we expected alterations in ICC firing to be reflected in ABR_LDS_ amplitudes. There are a few potential confounding factors. One is that the auditory stimuli were presented with both ears open during the ICC extracellular recordings but with one ear plugged during the ABR_LDS_. Although both sets of recordings were made from both sides, the extracellular ICC recordings would have been driven by simultaneous ipsilateral and contralateral inputs, while the ABRs were less so. Another factor is that ABRs exhibit primarily the onset response in ICC, while the ICC recordings were analyzed across the entire duration of the response. Other ABR_LDS_ analysis methods may capture the ICC activity more accurately.

### Effect of sound exposure

Extracellular recordings in the ICC showed differences in LDS-induced changes of sound-driven spike rates between tinnitus, non-tinnitus, and control mice. This result was not reflected in the ABR_LDS_, which showed a significant difference between tinnitus and non-tinnitus mice but not between any sound-exposed cohort and control mice. Control mice exhibited an intermediate suppression of ABR peak amplitude falling between tinnitus and non-tinnitus mice, and a study with a larger number of mice may result in a significant difference between sound-exposed and control.

With human subjects, there is no measure of previous exposure to loud sound. So, the lack of difference between control and sound-exposed conditions in the mice in our study may represent a problem when the ABR_LDS_ is used with human subjects. Unlike the controlled history of sound exposure in animal studies, tinnitus patients have an unknown noise exposure history and may not show a measurable hearing loss (Sharma et al. 2021, Waechter and Brannstrom 2015). Because different mechanisms may cause otological damage and tinnitus, the electrophysiological measures that can identify noise-induced tinnitus may not identify other types of tinnitus (Martines et al. 2010, Savastano 2008).

### Lateralization of tinnitus

In both the ICC and ABR_LDS_ recordings, the differences in afterdischarge activity and tone-evoked responses associated with tinnitus were often asymmetrical. Extracellular recordings showed tinnitus-specific differences for tone-driven firing in the contralateral ICC but not the ipsilateral side. However, both sides showed a tinnitus-specific difference in the influence of the LDS on tone-evoked and afterdischarge activity. With the ABR_LDS_, the delta ABR waveform amplitudes significantly differed between sound-exposed and non-exposed ears. These findings suggest that the neurological changes associated with tinnitus can be lateralized following sound exposure in the left ear alone.

Human tinnitus patients have reported unilateral tinnitus. However, it is hard to distinguish genuinely unilateral tinnitus from differences in interaural tinnitus levels that might merely cause the percept to seem lateralized. Imaging studies from human tinnitus patients demonstrate mixed evidence of lateralized neurological changes associated with tinnitus: functional magnetic resonance imaging (fMRI) of sound-evoked responses in the auditory cortex and IC showed less lateralization in human tinnitus patients than control patients when both groups have near-normal hearing (Lanting et al. 2014). In another fMRI study, patients with lateralized tinnitus exhibited abnormally low activation in the IC contralateral to the perceived tinnitus compared to patients with non-lateralized tinnitus (Melcher et al. 2000). Positron emission tomography (PET) imaging studies of patients with lateralized tinnitus have found abnormal activation contralateral to the perceived tinnitus in the auditory cortex (Lockwood et al. 1998), increased activity in the left side of the brain regardless of the perceived tinnitus (Mirz et al. 2000), and bilateral activation independent of tinnitus laterality (Giraud et al. 1999). Melcher et al., 2009, suggest that asymmetrical activation in the IC may be seen only in a subgroup of tinnitus patients, which may explain why multiple studies report different lateralization patterns (Melcher et al. 2009).

Evidence of lateralized neurological changes associated with tinnitus is similarly mixed in animal models. Rats with unilateral noise exposure demonstrate more tinnitus-like behavior when responding to acoustic stimuli presented to the sound-exposed ear (Heffner 2011). Another study found that unilaterally sound-exposed rats, not separated by behavioral evidence of tinnitus, did not exhibit significant differences in spontaneous firing rate between the contra- and ipsilateral ICs (Ropp et al. 2014).

### Frequency specificity and tinnitus

We found that tinnitus-specific differences in sound-evoked plasticity depended on the frequency of the probe tones, in ICC extracellular recordings and ABR_LDS_ recordings. ABR_LDS_ responses from non-tinnitus mice were significantly suppressed compared to tinnitus mice, but only when the stimulus frequency was at or above the sound exposure frequency. These results agree with human studies reporting that the tinnitus pitch corresponds with a region of impaired hearing (Henry and Meikle 1999, Norena et al. 2002, Sereda et al. 2011). There is also evidence that the tinnitus pitch is at the frequency of maximum hearing loss (Ochi et al. 2003), although this connection may only hold for a sub-group of tinnitus patients (Pan et al. 2009, Sereda et al. 2011). The finding that, with the ABR_LDS_, stimulating at frequencies above the sound exposure resulted in significant differences between tinnitus and non-tinnitus mice is consistent with previous studies showing frequency-specific changes in the auditory system with tinnitus.

### Hyperexcitability in the IC with tinnitus

Increased neuronal excitability in the IC has been established in animal models and human patients with tinnitus (Berger and Coomber 2015). Studies using fMRI imaging in human patients showed increased sound-driven activity in the IC of tinnitus subjects (Melcher et al. 2009). Elevated spontaneous activity (Mulders and Robertson 2013, Mulders and Robertson 2009) and increased burst firing (Bauer et al. 2008) have also been seen in the IC of animal models of tinnitus after noise exposure.

In this study, the data from tone-evoked ABR_LDS_ show a tinnitus-specific result in that the non-tinnitus mice had a more suppressed ABR_LDS_ amplitude after the LDS compared to tinnitus mice. It is unlikely to result from damage due to sound exposure because tinnitus mice were exposed to the same acoustic trauma as non-tinnitus mice. Therefore, we theorize that hyperactivity in the auditory brainstem in tinnitus mice compensates for reduced activity resulting from sound exposure. Because the lack of suppression in tinnitus mice was evident only in the late ABR waves, it is likely that more central auditory brainstem regions, such as the IC, are affected by the LDS in a tinnitus-specific manner. The LDS may not similarly affect sound-driven activity in more peripheral areas, such as the cochlear nucleus. This suggests that the IC may be essential for amplifying the relative increase in activity with tinnitus and that electrophysiological measurements of the IC could prove helpful in diagnosing tinnitus.

## 7 Author Contributions

EFS: Conceptualization, Formal analysis, Investigation, Methodology, Visualization, Writing-original draft, Writing-review and editing

CML: Data curation, Investigation, Software, Writing-editing and review

DLO: Conceptualization, Funding Acquisition, Resources, Supervision, Writing-review and editing

ALB: Conceptualization, Formal Analysis, Investigation, Methodology, Visualization, Writing-original draft, Writing-review and editing

## 9 Data Availability Statement

The raw data supporting the conclusions of this manuscript will be made available upon request, without undue reservation.

## Acknowledgements

A version of these data was included in Emily M. Fabrizio-Stover’s doctoral dissertation.

## Liturature Cited

Auerbach, B. D., Rodrigues, P. V. and Salvi, R. J. (2014) ‘Central gain control in tinnitus and hyperacusis’, Front Neurol, 5, 206.

Bauer, C. A., Turner, J. G., Caspary, D. M., Myers, K. S. and Brozoski, T. J. (2008) ‘Tinnitus and inferior colliculus activity in chinchillas related to three distinct patterns of cochlear trauma’, J Neurosci Res, 86(11), 2564–78.

Berger, J. I. and Coomber, B. (2015) ‘Tinnitus-related changes in the inferior colliculus’, Front Neurol, 6, 61.

Brozoski, T. J., Bauer, C. A. and Caspary, D. M. (2002) ‘Elevated fusiform cell activity in the dorsal cochlear nucleus of chinchillas with psychophysical evidence of tinnitus’, J Neurosci, 22(6), 2383–90.

Burghard, A. L., C; Fabrizio-Stover; E, Oliver, D (2022) ‘Long-Duration Sound-Induced Potentitation Changes Population Activity in the Inferior Colliculus’, Front Syst Neurosci.

Burghard, A. L., Lee, C. M., Fabrizio-Stover, E. M. and Oliver, D. L. (2022) ‘Long-Duration Sound-Induced Facilitation Changes Population Activity in the Inferior Colliculus’, Front Syst Neurosci, 16, 920642.

Burghard, A. L., Morel, N. P. and Oliver, D. L. (2019) ‘Mice heterozygous for the Cdh23/Ahl1 mutation show age-related deficits in auditory temporal processing’, Neurobiol Aging, 81, 47–57.

Cai, R., Montgomery, S. C., Graves, K. A., Caspary, D. M. and Cox, B. C. (2018) ‘The FBN rat model of aging: investigation of ABR waveforms and ribbon synapse changes’, Neurobiol Aging, 62, 53–63.

Cody, A. R. and Johnstone, B. M. (1981) ‘Acoustic trauma: single neuron basis for the “half-octave shift“’, J Acoust Soc Am, 70(3), 707–11.

Domarecka, E., Olze, H. and Szczepek, A. J. (2020) ‘Auditory Brainstem Responses (ABR) of Rats during Experimentally Induced Tinnitus: Literature Review’, Brain Sci, 10(12).

Eggermont, J. J. (2019) ‘Auditory brainstem response’, Handb Clin Neurol, 160, 451–464.

Fabrizio-Stover, E. M., Nichols, G., Corcoran, J., Jain, A., Burghard, A. L., Lee, C. M. and Oliver, D. L. (2022) ‘Comparison of two behavioral tests for tinnitus assessment in mice’, Front Behav Neurosci, 16, 995422.

Giraud, A. L., Chery-Croze, S., Fischer, G., Fischer, C., Vighetto, A., Gregoire, M. C., Lavenne, F. and Collet, L. (1999) ‘A selective imaging of tinnitus’, Neuroreport, 10(1), 1–5.

Heffner, H. E. (2011) ‘A two-choice sound localization procedure for detecting lateralized tinnitus in animals’, Behav Res Methods, 43(2), 577–89.

Henry, J. A. and Meikle, M. B. (1999) ‘Pulsed versus continuous tones for evaluating the loudness of tinnitus’, J Am Acad Audiol, 10(5), 261–72.

Holt, A. G., Bissig, D., Mirza, N., Rajah, G. and Berkowitz, B. (2010) ‘Evidence of key tinnitus-related brain regions documented by a unique combination of manganese-enhanced MRI and acoustic startle reflex testing’, PLoS One, 5(12), e14260.

Jacxsens, L., De Pauw, J., Cardon, E., van der Wal, A., Jacquemin, L., Gilles, A., Michiels, S., Van Rompaey, V., Lammers, M. J. W. and De Hertogh, W. (2022) ‘Brainstem evoked auditory potentials in tinnitus: A best-evidence synthesis and meta-analysis’, Front Neurol, 13, 941876.

Kalappa, B. I., Brozoski, T. J., Turner, J. G. and Caspary, D. M. (2014) ‘Single unit hyperactivity and bursting in the auditory thalamus of awake rats directly correlates with behavioural evidence of tinnitus’, J Physiol, 592(22), 5065–78.

Kaltenbach, J. A., Zhang, J. and Finlayson, P. (2005) ‘Tinnitus as a plastic phenomenon and its possible neural underpinnings in the dorsal cochlear nucleus’, Hear Res, 206(1-2), 200–26.

Kehrle, H. M., Granjeiro, R. C., Sampaio, A. L., Bezerra, R., Almeida, V. F. and Oliveira, C. A. (2008) ‘Comparison of auditory brainstem response results in normal-hearing patients with and without tinnitus’, Arch Otolaryngol Head Neck Surg, 134(6), 647–51.

Land, R., Burghard, A. and Kral, A. (2016) ‘The contribution of inferior colliculus activity to the auditory brainstem response (ABR) in mice’, Hear Res, 341, 109–118.

Lanting, C. P., de Kleine, E., Langers, D. R. and van Dijk, P. (2014) ‘Unilateral tinnitus: changes in connectivity and response lateralization measured with FMRI’, PLoS One, 9(10), e110704.

Lockwood, A. H., Salvi, R. J., Coad, M. L., Towsley, M. L., Wack, D. S. and Murphy, B. W. (1998) ‘The functional neuroanatomy of tinnitus: evidence for limbic system links and neural plasticity’, Neurology, 50(1), 114–20.

Longenecker, R. J. and Galazyuk, A. V. (2011) ‘Development of tinnitus in CBA/CaJ mice following sound exposure’, J Assoc Res Otolaryngol, 12(5), 647–58.

Martines, F., Bentivegna, D., Martines, E., Sciacca, V. and Martinciglio, G. (2010) ‘Characteristics of tinnitus with or without hearing loss: clinical observations in Sicilian tinnitus patients’, Auris Nasus Larynx, 37(6), 685–93.

McFadden, D. (1986) ’The curious half-octave shift: Evidence for basalward migration of the traveling-wave envelope with increasing intensity’ in Richard J. Salvi, D. H. R.P. Hamernik, V. Colletti, ed. Basic and Applied Aspects of Noise-Induced Hearing Loss, Plenum publishing corporation, 295–312.

Melcher, J. R. and Kiang, N. Y. (1996) ‘Generators of the brainstem auditory evoked potential in cat. III: Identified cell populations’, Hear Res, 93(1-2), 52–71.

Melcher, J. R., Levine, R. A., Bergevin, C. and Norris, B. (2009) ‘The auditory midbrain of people with tinnitus: abnormal sound-evoked activity revisited’, Hear Res, 257(1-2), 63–74.

Melcher, J. R., Sigalovsky, I. S., Guinan, J. J., Jr. and Levine, R. A. (2000) ‘Lateralized tinnitus studied with functional magnetic resonance imaging: abnormal inferior colliculus activation’, J Neurophysiol, 83(2), 1058–72.

Milloy, V., Fournier, P., Benoit, D., Norena, A. and Koravand, A. (2017) ‘Auditory Brainstem Responses in Tinnitus: A Review of Who, How, and What?’, Front Aging Neurosci, 9, 237.

Mirz, F., Gjedde, A., Ishizu, K. and Pedersen, C. B. (2000) ‘Cortical networks subserving the perception of tinnitus--a PET study’, Acta Otolaryngol Suppl, 543, 241–3.

Mohrle, D., Ni, K., Varakina, K., Bing, D., Lee, S. C., Zimmermann, U., Knipper, M. and Ruttiger, L. (2016) ‘Loss of auditory sensitivity from inner hair cell synaptopathy can be centrally compensated in the young but not old brain’, Neurobiol Aging, 44, 173–184.

Mulders, W. H. and Robertson, D. (2009) ‘Hyperactivity in the auditory midbrain after acoustic trauma: dependence on cochlear activity’, Neuroscience, 164(2), 733–46.

Mulders, W. H. and Robertson, D. (2013) ‘Development of hyperactivity after acoustic trauma in the guinea pig inferior colliculus’, Hear Res, 298, 104–8.

Norena, A., Micheyl, C., Chery-Croze, S. and Collet, L. (2002) ‘Psychoacoustic characterization of the tinnitus spectrum: implications for the underlying mechanisms of tinnitus’, Audiol Neurootol, 7(6), 358–69.

Ochi, K., Ohashi, T. and Kenmochi, M. (2003) ‘Hearing impairment and tinnitus pitch in patients with unilateral tinnitus: comparison of sudden hearing loss and chronic tinnitus’, Laryngoscope, 113(3), 427–31.

Ono, M., Bishop, D. C. and Oliver, D. L. (2016) ‘Long-Lasting Sound-Evoked Afterdischarge in the Auditory Midbrain’, Sci Rep, 6, 20757.

Pan, T., Tyler, R. S., Ji, H., Coelho, C., Gehringer, A. K. and Gogel, S. A. (2009) ‘The relationship between tinnitus pitch and the audiogram’, Int J Audiol, 48(5), 277–94.

Parthasarathy, A. and Kujawa, S. G. (2018) ‘Synaptopathy in the Aging Cochlea: Characterizing Early-Neural Deficits in Auditory Temporal Envelope Processing’, J Neurosci, 38(32), 7108–7119.

Ropp, T. J., Tiedemann, K. L., Young, E. D. and May, B. J. (2014) ‘Effects of unilateral acoustic trauma on tinnitus-related spontaneous activity in the inferior colliculus’, J Assoc Res Otolaryngol, 15(6), 1007–22.

Sametsky, E. A., Turner, J. G., Larsen, D., Ling, L. and Caspary, D. M. (2015) ‘Enhanced GABAA-Mediated Tonic Inhibition in Auditory Thalamus of Rats with Behavioral Evidence of Tinnitus’, J Neurosci, 35(25), 9369–80.

Savastano, M. (2008) ‘Tinnitus with or without hearing loss: are its characteristics different?’, Eur Arch Otorhinolaryngol, 265(11), 1295–300.

Sereda, M., Hall, D. A., Bosnyak, D. J., Edmondson-Jones, M., Roberts, L. E., Adjamian, P. and Palmer, A. R. (2011) ‘Re-examining the relationship between audiometric profile and tinnitus pitch’, Int J Audiol, 50(5), 303–12.

Sharma, A., Sood, N., Munjal, S. and Panda, N. (2021) ‘Perception of Tinnitus Handicap And Stress Across Age Groups in Normal Hearing’, Int Tinnitus J, 25(1), 13–17.

Singh, S., Munjal, S. K. and Panda, N. K. (2011) ‘Comparison of auditory electrophysiological responses in normal-hearing patients with and without tinnitus’, J Laryngol Otol, 125(7), 668–72.

Waechter, S. and Brannstrom, K. J. (2015) ‘The impact of tinnitus on cognitive performance in normal-hearing individuals’, Int J Audiol, 54(11), 845–51.

